# Inhibition tunes prefrontal circuit dynamics to promote sociosexual behavior in female mice

**DOI:** 10.1101/2025.08.06.668891

**Authors:** Elizabeth A. Amadei, Pau Vilimelis Aceituno, Reinhard Loidl, Roman Boehringer, Eduarda Streit Morsch, Benjamin Ehret, Benjamin F. Grewe

## Abstract

The prefrontal cortex (PFC) plays a central role in the selection and expression of diverse social behaviors. A balance of excitation and inhibition is necessary for normal social functioning, but it remains unclear how inhibition sculpts pyramidal activity to promote specific social behaviors. To address this question, we developed a novel decision-making task in which female mice chose between interaction with a male (sociosexual stimulus) or an appetitive non-social option (milk solution). To explore the role of a task-relevant inhibitory subpopulation, we targeted neurons in the medial PFC (mPFC) expressing oxytocin receptors (OXTR neurons). Combining optogenetic inhibition of OXTR neurons with population calcium imaging of pyramidal neural activity, we found that OXTR neurons normally promote interaction with a male compared to a non-social alternative. OXTR neurons also regulate pyramidal activity, which enables a specific ensemble to represent the male option during decision-making. Computational modeling reproduced these findings through a pyramidal competition mechanism, in which regulation by OXTR neurons allows a male-representing pyramidal ensemble to effectively compete against the remaining pyramidal population to drive male choice. These results provide a candidate mechanism by which inhibition enables mPFC pyramidal activity to select for specific social behaviors and may help explain why excitation/inhibition balance is so important for social functioning.

## Introduction

Natural environments are rich with social and non-social behavioral options, which each confer important benefits. While social interactions are important for reproduction, resource exchange, and the formation and maintenance of relationships, non-social behaviors (e.g., foraging, sleeping) are critical for self-preservation^1^. Prioritizing and effectively engaging in social behaviors is essential for mental health, with disorders such as autism spectrum disorder, Williams syndrome, schizophrenia, and anxiety featuring social deficits^2–5^.

The prefrontal cortex (PFC) plays a key role in social behavioral selection and expression^6–8^. In general, PFC is thought to integrate internal (e.g., state, memory) and external (e.g., stimuli) variables to guide ongoing behavior^9^. In social contexts, these variables include both self- and other-related information such as hormonal and affective states, history of interaction, sex, age, and group hierarchy^7^. Through the PFC and its projections, these variables are translated into behaviors such as social preference or avoidance^10–12^, discrimination of others’ affective states^13^, mating^14^, aggression^15^, and dominance or submission behaviors^16,17^.

Rodent studies on the medial PFC (mPFC) have shown that normal social functioning requires a balance of neural excitation and inhibition^18–20^. For example, artificially elevating mPFC pyramidal activity disrupted mice’ preference for a novel conspecific (over a neutral chamber), but this preference could be restored by simultaneously activating parvalbumin-expressing (PV) interneurons^18^. Further, boosting PV neurons’ activity or dampening pyramidal activity rescued social interaction in a *CNTNAP2*-knockout mouse model of autism, suggesting that reduced inhibitory capacity may contribute to social dysfunction in this disorder^20,21^.

In addition to PV neurons, other interneuron types in mPFC play a key role in social functioning. For example, mPFC somatostatin (SST) interneurons are necessary for affective state discrimination, where mice prefer to spend time with a conspecific in an altered affective state (e.g., stressed, relieved) compared to a neutral state^13^. mPFC neurons expressing receptors for the neurochemical oxytocin (OXTR) are necessary for sociosexual behavior in female mice. Inhibiting these neurons disrupted females’ preference for a male compared to a novel object during the sexually-receptive (estrus) phase of the female’s estrous cycle^22^. While there is some evidence for mPFC pyramidal neurons expressing OXTR^23,24^, OXTR neurons are predominately a subpopulation of interneurons (putatively SST)^22^, which locally inhibit pyramidal neurons^24,25^. Consistently, inhibiting the full population of SST interneurons had the same effect as inhibiting OXTR neurons on reducing subjects’ male preference, whereas inhibiting pyramidal neurons had no effect^22^. These results suggest that mPFC OXTR interneurons support female sociosexual behavior.

Based on this work, one fundamental open question is how interneurons promote specific social behaviors over other social or non-social options. One possibility is that they bias a competition between pyramidal cell groups representing these behavioral options, amplifying the activity of one ensemble over others^8^. This difference in activity could then be read out by downstream areas to execute a behavioral choice^8^. However, this specific possibility remains untested.

To address this question, we designed a novel decision-making task in which female mice chose between a sociosexual option (male stimulus) or an appetitive non-social alternative (milk solution) on many trials. We then asked how a task-relevant inhibitory source (mPFC OXTR neurons) regulates pyramidal activity and subjects’ ability to choose the male option. We combined optogenetic inhibition of mPFC OXTR neurons with imaging of pyramidal population activity during this task.

We found that inhibiting OXTR neurons reduced subjects’ behavioral choice of the male, increased pyramidal population activity and disrupted the ability of a specific pyramidal ensemble to represent the male option during decision-making. Computational modeling reproduced our experimental findings through a pyramidal competition mechanism, in which OXTR neurons regulate the ability of a male-representing pyramidal ensemble to drive male choice. Our results reveal a potential circuit mechanism by which inhibition sculpts prefrontal pyramidal activity to promote specific social behaviors.

## Results

### Inhibiting mPFC OXTR neurons while imaging pyramidal activity

To explore the role of mPFC OXTR neurons in sociosexual decision-making, we expressed Cre-dependent, mCherry-tagged Halorhodopsin or an inactive control fluorophore in the right mPFC of heterozygous OXTR-Cre female mice (*n* = 8 Halo subjects, *n* = 8 control subjects; Figures 1A, S1). This allowed us to optogenetically inhibit OXTR neurons, which we verified using slice electrophysiology (Figures S2-3). To simultaneously image mPFC pyramidal population activity, we co-expressed the fluorescent calcium indicator GCaMP6m under control of the CaMKIIα promoter (Figures S1A-E). To deliver light for both optogenetic inhibition and population calcium imaging, we implanted a gradient-index lens above the right mPFC (Figures S1A, F), which interfaced with a miniaturized microscope system. This approach allowed us to combine optogenetic inhibition of OXTR neurons with imaging of pyramidal population activity in freely behaving subjects.

**Figure 1.**
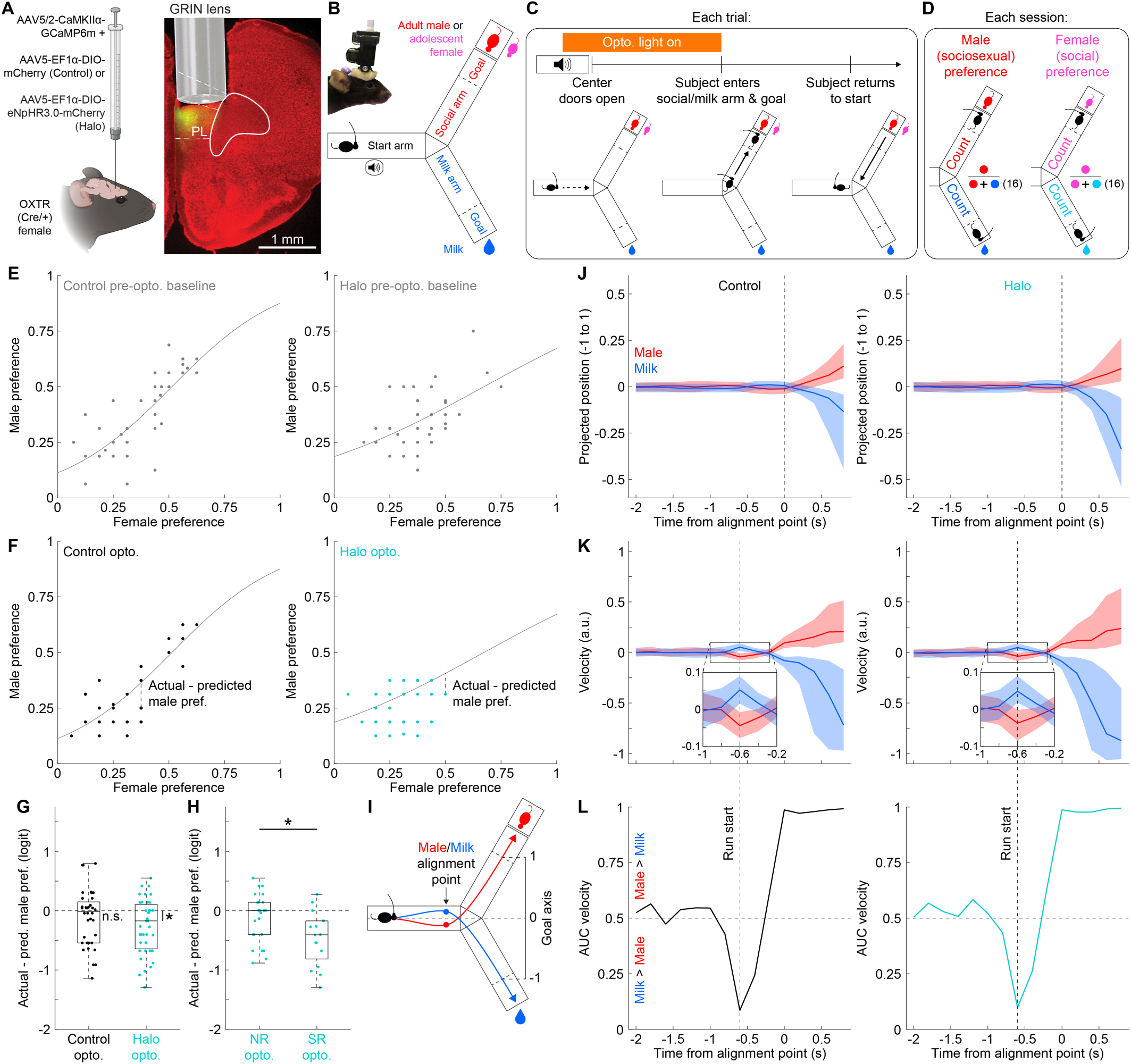
Inhibiting mPFC OXTR neurons in female mice decreases male choice in a task of sociosexual decision-making. (**A**) Experimental groups virally expressing imaging and optogenetic constructs in the right mPFC. Female Cre-expressing mice expressed GCaMP6m in pyramidal neurons and either Halorhodopsin (Halo) or a control fluorophore (control) in OXTR neurons (*n* = 8, 7 respectively). One additional fluorophore-injected subject was Cre-negative and was therefore included in the control group (*n* = 8 total). A GRIN lens was implanted directly above the expression site for optical imaging and optogenetic light delivery. PL: prelimbic cortex. (**B**) Behavioral task, in which freely-moving subjects chose between social interaction or a sweetened milk solution on each trial. Within a given session, subjects were exposed to both a novel adult male and adolescent female (each for half the total number of trials, typically 16 out of 32). (**C**) Trial structure, including timing of optogenetic light delivery. (**D**) Calculation of male and female behavioral preference in each session. Male preference is the proportion of trials that the subject chose the male stimulus while he was present in the arena (typically 16 trials). Female preference is the proportion of trials that the subject chose the adolescent female stimulus while she was present in the arena (typically 16 trials). (**E**) Female versus male preference over control (left, *n* = 40) and Halo (right, *n* = 42) sessions before the introduction of optogenetic light (pre-opto. baseline sessions). A binomial logistic regression model was fit to each dataset (overlaid curves; fixed effect of female preference (mean-centered); control: *Estimate* = 4.007, *SE* = 0.610, *Z* = 6.57, *P* < 0.001***; Halo: *Estimate* = 2.192, *SE* = 0.646, *Z* = 3.40, *P* < 0.001***). (**F**) Female versus male preference over control (left, *n* = 36) and Halo (right, *n* = 39) sessions with optogenetic light delivery (opto. sessions). Overlaid are baseline fits from (**E**), which were used to test whether male preference during opto. differed from baseline (actual male preference during opto. versus predicted male preference from baseline fit). (**G**) Actual - predicted male preference in control and Halo opto. sessions (same *n* as (**F**)). Session values transformed to logit scale to match binomial generalized linear mixed model (GLMM) used for statistical testing. Boxplots show median and interquartile range (IQR) over sessions. Only Halo group showed a significant difference between actual and predicted male preference (control: *Estimate* = -0.097, *SE* = 0.110, *Z* = -0.88, *P* = 0.380; Halo: *Estimate* = -0.225, *SE* = 0.107, *Z* = -2.11, *P* = 0.035*). In control group only, actual and predicted male preference were significantly equivalent within a tolerance range of +/- 1 male trials (Two One-Sided Tests (TOST) procedure: *P control* = 0.025*, *P Halo* = 0.213). (**H**) Actual - predicted male preference on non-receptive (NR, *n* = 23) and sexually receptive (SR, *n* = 16) Halo opto. sessions. Boxplots show median and IQR over sessions. Estrous state (SR vs. NR) modulated the difference between actual and predicted male preference (binomial GLMM with estrous state as fixed effect; *Estimate* = -0.329, *SE* = 0.190, *Z* = -1.73, *P (one-sided)* = 0.042*). (**I**) On each trial, position was projected onto goal axis and aligned to start of directed movement towards male or milk goal. (**J-K**) Projected position (**J**) and corresponding velocity (**K**) relative to alignment point on male and milk trials in control (left) and Halo (right) opto. sessions. Thick lines and shaded areas show median and IQR over pooled trials (control: *n* = 173 male, 402 milk; Halo: *n* = 162, 461). (**L**) AUC of velocity on male versus milk trials from (**K**). Vertical dotted line indicates run start (0.6 s before alignment point).

### Sociosexual decision-making task

We then trained subjects to perform a novel decision-making task, in which they chose between social interaction or drinking of a sweetened milk solution on each trial (Figure 1B; full experiment timeline in Figure S4A and Methods). The social option consisted of a novel conspecific in a “social chamber”, which had one side partially open (vertical bars) to allow for social (sniffing) interactions. The milk option consisted of a droplet of milk solution dispensed from a syringe system. Subjects underwent water scheduling to ensure motivation to drink the milk solution (see Figure S4A and Methods).

Each trial was initiated by a 5 s, 4 kHz tone. Then, doors at the center of the arena opened, allowing the subject to enter either the social or milk arm. Once the subject reached the social or milk goal, it had 15 s to socially interact or drink a milk droplet, before a door in front of the goal closed, and the subject had to return to the start arm to initiate the next trial (Figures 1C, S4B). Each daily session consisted of up to 32 trials.

Given that both sociosexual motivation and general social interest could contribute to the subject choosing a male stimulus in this task, we aimed to isolate the effect of sociosexual motivation by including a non-sexual social stimulus as well. We therefore exposed female subjects to two types of social stimuli in each session, each for half of the total number of trials (typically 16 out of 32). These stimuli were an adult male (sociosexual stimulus; P63-112) and an adolescent female (non-sexual social stimulus; P25-38)^26^. The female subject began the session in the presence of one of the stimuli (e.g., male), choosing between that stimulus and the milk option. Once the subject completed half the total number of trials, the social stimulus was replaced (e.g., male stimulus removed, and female stimulus placed in arena). The social presentation order (male or female first), as well as the locations of the social and milk options (left or right relative to start arm), were fixed over sessions within a given subject, but varied over subjects (Figure S4C). All combinations of presentation order and social/milk location were represented in both control and Halo groups (Figure S4C). Importantly, including a non-sexual social stimulus allowed us to isolate the effect of inhibiting OXTR neurons on sociosexual decision-making, consistent with previous work on sociosexual motivation^22^.

To determine whether subjects’ hormonal state interacted with inhibition of OXTR neurons to affect decision-making, we tracked the estrous cycle of each subject on each session using vaginal cytology^27–29^ (Figure S4D). We classified each session as being either putatively sexually receptive (SR; proestrus and early estrus phases of estrous cycle) or non-receptive (NR; all other estrous phases; see Figure S4D and Methods)^30,31^.

Once subjects learned and stably performed the task (Figures S4E-G), we introduced optogenetic light stimulation to inhibit mPFC OXTR neurons (starting from session 6 or 7; optogenetic (opto.) sessions). To influence decision-making, we delivered light automatically from the second half of the tone to the moment the subject entered either the social or milk arm on every trial (Figure 1C; Videos S1-2). The Halo and control groups received similar light exposure (Figure S4H) and showed a similar weight drop from water scheduling (Figure S4I), confirming a matched experimental design.

### Inhibiting OXTR neurons reduces male choice

To determine whether and how inhibiting OXTR neurons affected sociosexual decision-making, we took all trials where the male was present in each session (up to 16) and computed the proportion of trials that the subject chose the male (“male preference”, Figure 1D). As a non-sexual social comparison, we performed the same analysis on trials where the female was present in the session (up to 16), computing the female preference (Figure 1D). This gave two preference values per session (male and female preference).

We then established a baseline relationship between male and female preference by taking these values in early decision-making sessions before the introduction of optogenetic stimulation on day 6 or 7 (pre-opto. sessions, Figures S4A, F-G). Plotting male preference relative to female preference over sessions (Figure 1E) revealed a significant relationship in both control and Halo groups (shown for each group in Figure 1E as fit of binomial logistic regression model). The two groups were well-matched in that they showed a similar distribution of the predictor variable (female preference, Figure S5A) and no systematic bias from their fitted curve (Figure S5B).

Establishing a baseline fit between male and female preference allowed us to test whether introducing optogenetic stimulation affected sociosexual decision-making. This effect would appear as a vertical shift of opto. sessions below the pre-opto. baseline curve (i.e., reduction in male preference; Figure 1F). To test this idea, we plotted female versus male preference in opto. sessions, and compared these values to the pre-opto. curve (Figure 1F).

As expected, opto. sessions in the control group consistently followed the pre-opto. baseline curve, such that there was no significant difference between actual male preference and that predicted using the baseline curve (Figure 1G). Further, the actual and predicted male preference were equivalent to each other within an expected chance-level tolerance range of +/- 1 male trials (Figure 1G). In contrast, opto. sessions in the Halo group showed a significant shift downward, such that male preference was significantly lower than predicted (Figure 1G). The absence of a shift in the control group and presence of a shift in the Halo group was not simply explained by a different distribution of female preference in opto. sessions, as female preference was similar between groups (Figure S5C). These results suggest that inhibiting OXTR neurons disrupts male choice.

### Inhibiting OXTR neurons interacts with hormonal state to reduce male choice

Based on previous work implicating mPFC OXTR neurons in female sociosexual motivation during sexual receptivity^22^, we hypothesized that inhibiting OXTR neurons would disrupt male choice most strongly during sexual receptivity. We therefore compared the shift in male preference (actual - predicted) between SR and NR opto. sessions in the Halo group (Figure 1H). We found a larger shift on SR sessions, suggesting that inhibiting OXTR neurons disrupts male choice most strongly during sexual receptivity.

Together, our results suggest that mPFC OXTR neurons in female mice promote male choice in the presence of an appetitive non-social option, most prominently during sexual receptivity.

### Characterizing the timing of behavioral choice on each trial

One key question arising from these findings was how choice unfolded within each trial. To detect choice, we tracked subjects’ center-of-mass position (XY pixel coordinates) as they ran to the male or milk goal on each trial. We aimed to identify the start of the run as a clear timestamp of subjects’ behavioral choice, to which we could later align neural activity. We first projected the position at each time sample onto an axis connecting the goal entry locations (Figures 1I-J). This allowed us to identify the timepoint at which subjects began monotonically moving toward their intended goal on each trial, which we referred to as the “alignment point” (Figures 1I-J). In both the control and Halo groups, subjects showed an interesting velocity pattern before the alignment point (Figure 1K). Specifically, subjects would first move further away from their intended goal (Figure 1K, zoom-in) before curving into a direct trajectory toward the goal at the alignment point. By comparing the velocity on male versus milk trials using Receiver-Operating Characteristic Curve (ROC) analysis, we found that velocity first strongly deviated between trial types at 0.6 seconds before the alignment point (prominent (negative) peak in area under the curve (AUC); Figure 1L). We therefore defined the run start on each trial at this timepoint.

### Inhibiting OXTR neurons increases pyramidal population activity

During optogenetic inhibition of mPFC OXTR neurons, we simultaneously performed population calcium imaging of mPFC pyramidal neurons (Figures 2A-B). This allowed us to measure how inhibiting OXTR neurons during decision-making affected pyramidal activity. We started by characterizing the change in activity in individual pyramidal neurons immediately when the optogenetic light turned on in each trial (first 2.5 s of light, Figure 2C). Within each session, we computed the proportion of neurons that increased, decreased or showed no change in activity. The Halo group showed a larger proportion of modulated neurons (increased or decreased activity) compared to control (Figure 2C, right).

**Figure 2.**
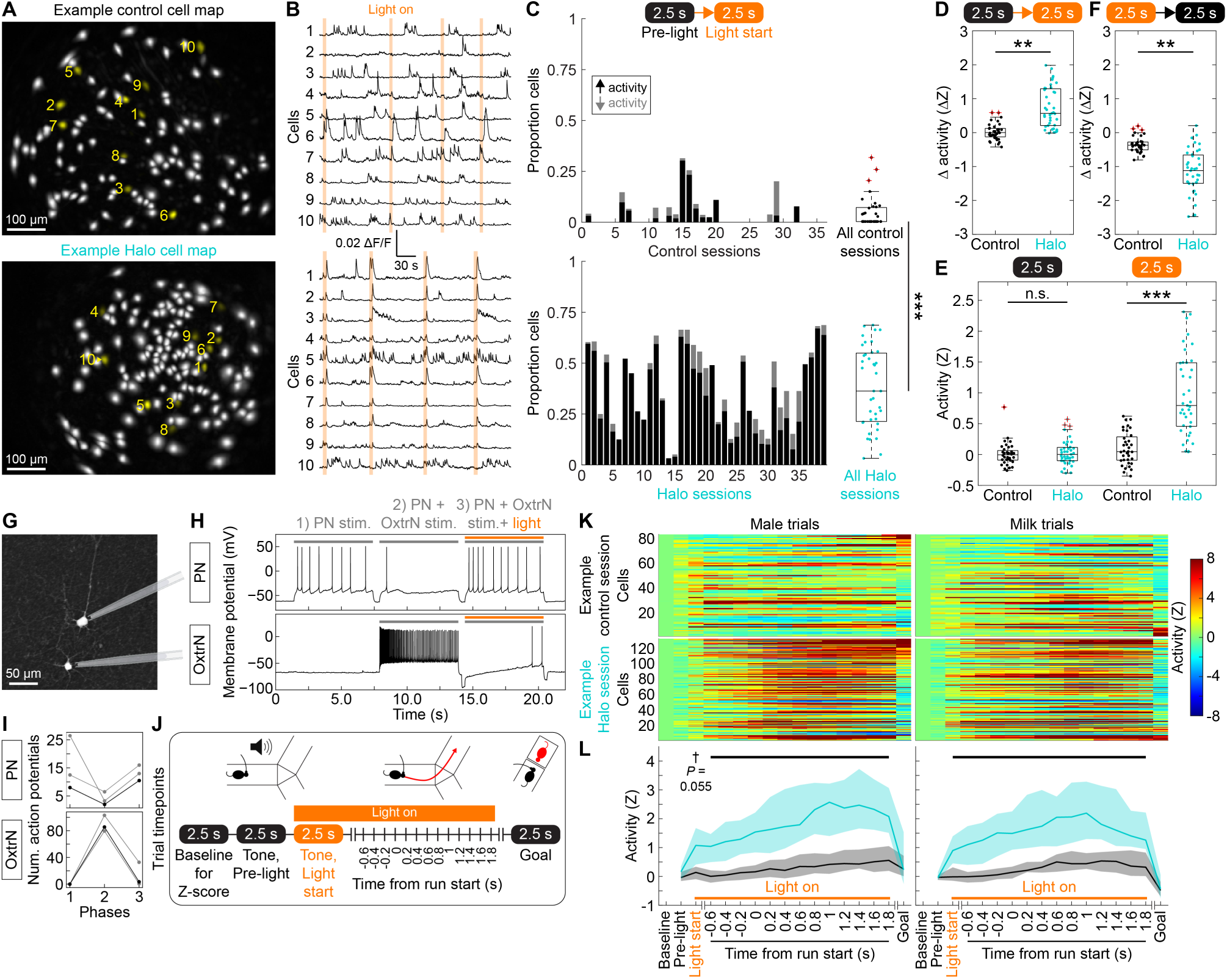
Inhibiting mPFC OXTR neurons increases pyramidal population activity. (**A**) Cell maps in example control and Halo sessions (*n* = 99 and 131 cells, respectively; median of 101 cells per session over all sessions). (**B**) Activity traces of 10 example cells highlighted in (**A**). Optogenetic light overlaid in orange. (**C**) Proportions of cells in each session that significantly increased (black) or decreased (grey) their activity in response to optogenetic light turning on (Wilcoxon signed-rank test over male and milk trials in each session, *P* < 0.05 after Benjamini-Hochberg correction; *n* = 36 control, 39 Halo sessions here and (**D-F, L**)). All remaining cells (not shown) showed no significant change. Right: Proportions of modulated cells (increased or decreased activity) in control versus Halo sessions (same *n* as Left). Halo group showed higher proportion of modulated cells (binomial GLMM with group as fixed effect: *Estimate* = 3.101, *SE* = 0.418, *Z* = 7.43, *P* < 0.001***). (**D**) Population response to optogenetic light turning on (male and milk trials) in control and Halo sessions. Halo group showed stronger boost of population activity compared to control (LME with group as fixed effect: *Estimate* = 0.745, *SE* = 0.183, *t*(13.95) = 4.07, *P* = 0.001**). (**E**) Population activity immediately before (left) and at the start (right) of optogenetic light in control and Halo sessions (LME with group (control, Halo), and timepoint (pre-light, light start) as fixed effects; significant group x timepoint interaction (*F*(1, 132.11) = 51.87, *P* < 0.001***). Groups only differed at the start of optogenetic light (pre-light (control - Halo): *Estimate* = -0.040, *SE* = 0.112, *t*(25.8) = -0.35, *P* = 0.727; light start (control - Halo): *Estimate* = -0.884, *SE* = 0.112, *t*(25.8) = -7.87, *P* < 0.001***; Holm-corrected *P* values). (**F**) Population response to optogenetic light turning off (male and milk trials) in control and Halo sessions. Halo group showed stronger drop of population activity compared to control (LME with group as fixed effect: *Estimate* = -0.815, *SE* = 0.238, *t*(13.99) = -3.43, *P* = 0.004**). (**G**) Representative reconstructed pair of pyramidal (PN; top) and Halorhodopsin-expressing OXTR (OxtrN; bottom) neurons in L2/3 mPFC. Both neurons were fluorescently labeled and reconstructed using biocytin-streptavidin staining. (**H**) Membrane potential recordings of neuron pair in (**G**) during three 6-s long experimental phases. (**I**) Number of action potentials in each phase for 3 pairs (Top: PN, Bottom: OxtrN). Pair from (**G-H**) shown in black. (**J**) Trial timepoints, which were either individual time samples (vertical lines) or windows containing multiple time samples (bubbles). Within windows, activity was averaged over samples to obtain one value per window. Orange bar indicates optogenetic light, which could extend through 1.8 s (from run start) on at least some trials (Figure S6B). (**K-L**) Neural activity over trial timepoints in two example sessions (**K**) and all sessions (**L**). Activity plotted separately on male (left) and milk (right) trials. (**K**) Activity of individual cells (rows) in example sessions, sorted by difference in activity between male and milk trials at goal timepoint. (**L**) Comparison of population activity between all control and Halo sessions over time. Compared to control group, population activity in Halo group was significantly elevated at timepoints with optogenetic light. The only exception was the “Tone, Light start” timepoint on male trials, where the *P* value was trending for significance after correcting for multiple comparisons (LME with group (control, Halo), timepoint, and trial type (male, milk) as fixed effects; significant group x timepoint x trial type interaction (*F*(15, 2190) = 2.51, *P* = 0.001**), followed by post hoc group comparisons at each timepoint; black bar indicates *P* < 0.05 after Holm correction for multiple comparisons). Throughout the figure, boxplots and thick lines/shaded areas show median and IQR over sessions.

To assess the overall effect on population activity, we summarized the change in activity over all cells in each session (Figure 2D). Compared to the control group, the Halo group showed a significant increase in population activity. This increase was not simply due to a difference in pre-light activity, as pre-light activity was comparable between groups (Figure 2E, left). Instead, it was due to elevated activity in the Halo group at the light start (Figure 2E, right). Elevated activity at the light start was not simply explained by movement, as there was no association between population activity and movement speed at this timepoint over sessions (linear mixed-effects model (LME) with mean-centered speed as fixed effect: *Estimate* = -0.084, *SE* = 0.067, *t*(71.65) = -1.25, *P* = 0.216). Finally, once the optogenetic light turned off, pyramidal population activity decreased in the Halo group compared to control (Figure 2F), consistent with a light-induced increase in pyramidal activity. Together, these results suggest that mPFC OXTR neurons normally regulate pyramidal population activity.

### Monosynaptic regulation of pyramidal activity by OXTR neurons

To explore the mechanism by which OXTR neurons regulate pyramidal activity, we performed dual whole-cell current-clamp recordings in acute brain slices of mPFC. We recorded from pyramidal and mCherry-tagged, Halorhodopsin-expressing OXTR neurons (Figures 2G-I). Previous work has shown that mPFC OXTR neurons are a subset of SST interneurons^22^, and both OXTR neurons and SST interneurons form monosynaptic connections onto pyramidal neurons^24,32^. We therefore hypothesized that increased pyramidal activity resulting from OXTR neuronal inhibition would occur in monosynaptically-connected pairs of pyramidal and OXTR neurons.

We identified pyramidal and OXTR neurons in slice based on cell morphology and mCherry expression, respectively. We identified putative monosynaptic pairs of pyramidal and OXTR neurons by voltage-clamping the pyramidal neuron at a depolarized value (∼40 mV), stimulating the OXTR neuron, and confirming current responses in the pyramidal neuron^32^. We then switched to current-clamp recordings and stimulated the pyramidal neuron (1) alone, (2) while stimulating the OXTR neuron, and (3) while stimulating the OXTR neuron in the presence of inhibitory optogenetic light (Figure 2H). In all three pairs of recorded neurons, stimulating the OXTR neuron reduced action potential firing in the pyramidal neuron (relative to its “alone” phase (phase 2 compared to 1); Figure 2I). However, turning on optogenetic light to inhibit the OXTR neuron was sufficient to increase pyramidal activity (Figure 2I). This single-cell finding paralleled our population-level finding that inhibiting OXTR neurons increased pyramidal population activity. This suggests that monosynaptic connectivity may explain, at least in part, the normal regulation of pyramidal population activity by OXTR neurons.

### Optogenetic inhibition of OXTR neurons and corresponding increase in pyramidal population activity coincides with decision-making

In addition to analyzing neural activity around the start or end of optogenetic light delivery, we tracked activity over the course of each trial, from the trial-start tone to when the subject reached the male or milk goal. We therefore extracted each cell’s activity during key timepoints within a trial, including the tone, run start, and arrival at the goal zone (Figure 2J-K). These timepoints were defined to be largely non-overlapping (Figure S6A). In contrast, the optogenetic light partially overlapped with these timepoints, spanning from the second half of the tone (“Tone, Light start” timepoint in Figure 2J) to the subject’s entrance in the social or milk arm (early part of the run). Given that trials varied in how long subjects took to complete this part of the run, the duration of optogenetic light and its overlap with run-associated timepoints also varied over trials. Specifically, all male and milk trials had optogenetic light through 0.8 seconds into the run (Figure S6B). Then, depending on running speed, the proportion of trials with optogenetic light decreased over subsequent timepoints. Optogenetic light was off on all trials by the goal timepoint (Figures 2J, S6B).

Comparing pyramidal population activity between Halo and control groups over timepoints, the Halo group showed elevated activity on timepoints with optogenetic light (Figures 2K-L). The only exception was the second half of the tone on male trials (“Tone, Light start” timepoint), which was trending for significance (*P* = 0.055) after correcting for multiple comparisons. These results indicate that inhibiting OXTR neurons elevates pyramidal population activity at behaviorally-relevant timepoints, including during putative decision-making (pre-run timepoints). These results also led us to ask how elevated population activity could affect the computation of the decision and ability to choose the male option.

### Identifying male-representing (MALE) cells

mPFC pyramidal activity normally discriminates between social and non-social stimuli and rewards^33–35^. On the other hand, mouse models of autism and Rett syndrome show reduced neural discriminability and sociability^33,34^. Based on these findings, it has been proposed that the ability of mPFC to distinctly represent social stimuli is necessary for selecting and/or maintaining social behaviors^33,34^. Consistently, re-activating mPFC neurons responsive to either male or female stimuli has been shown to promote a behavioral bias for that stimulus type in a sex preference assay^11^. We therefore hypothesized that (1) neurons which preferentially represent the male stimulus in our task are recruited during decision-making to promote male choice and (2) elevated pyramidal population activity (resulting from inhibiting OXTR neurons) disrupts this representation and the ability to choose the male option.

We started this analysis in control sessions. To identify male-representing neurons in each session, we took each neuron’s activity at the goal timepoint on male and milk trials. We then asked how well the neuron discriminated between the two trial types in the session using Receiver Operating Characteristic (ROC) analysis (Figure 3A). We took the area under the ROC curve (AUC) and rescaled it between -1 and 1 (“separability score”)^36^, such that a positive value would indicate a male preference, a negative value would indicate a milk preference, and a value of 0 would indicate no preference. Neurons showed a range of separability values between -1 and 1 (Figure 3B).

**Figure 3.**
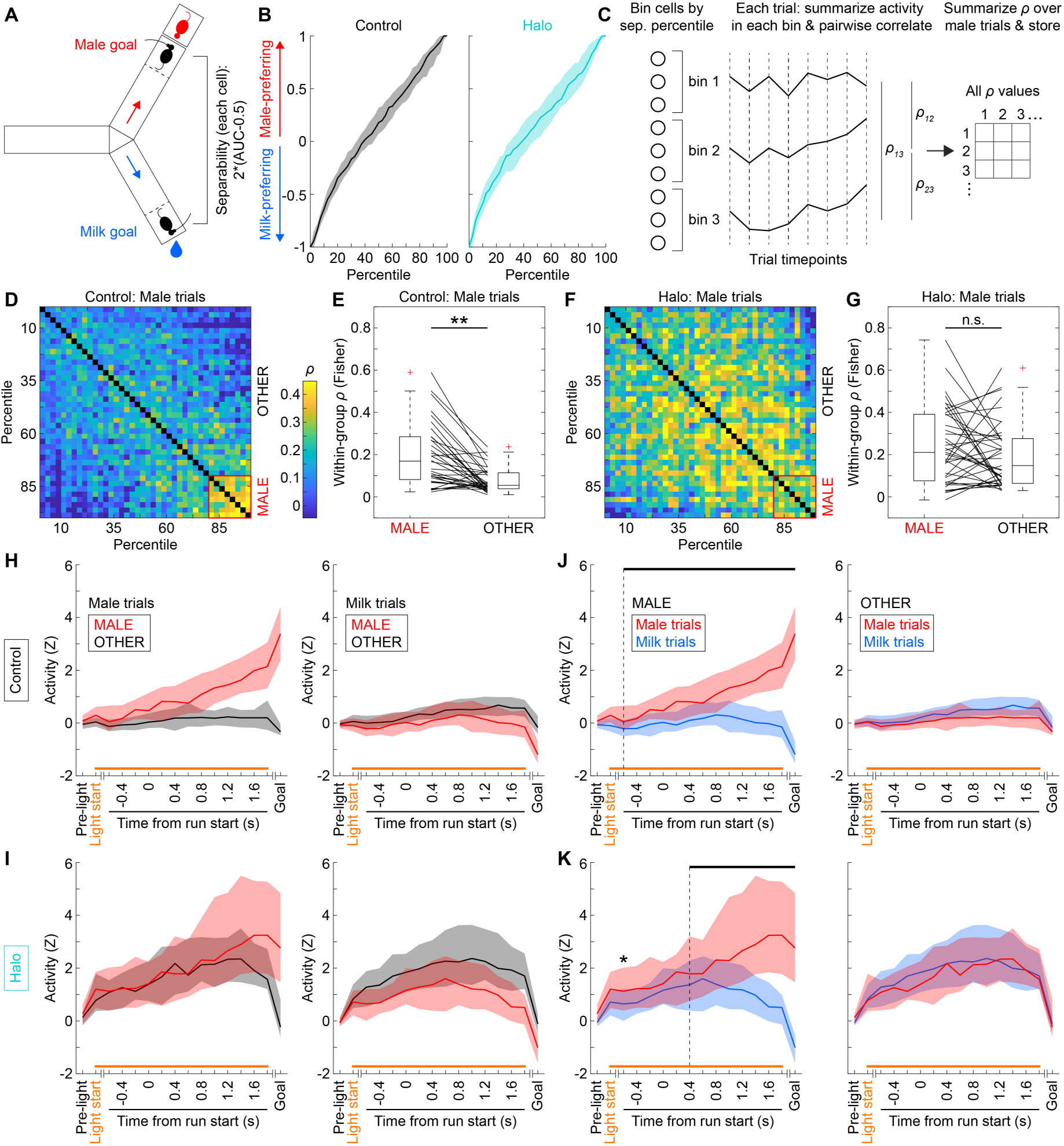
Inhibiting mPFC OXTR neurons delays the ability of a specific ensemble to represent the male option. (**A**) Definition of separability score. In each session, the ability of each cell to discriminate between male and milk trials was evaluated at the goal timepoint using ROC AUC analysis. Each cell’s AUC was rescaled between -1 to 1 to define a separability score. A positive value indicated preference for the male, a negative value indicated preference for the milk, and a value of 0 indicated no preference. (**B**) Distribution of separability values over cells in control (left, *n* = 36) and Halo (right, *n* **=** 39) sessions (same *n* throughout figure). (**C**) Approach to define male-representing (MALE) cells in each session, involving pairwise correlating the activity of cells that were sorted and binned by their separability value (2.5 percentile-width bins; 40 bins total). Activity was taken on male trials. This approach produced a matrix of Spearman correlation coefficients for each session. (**D**) Summary coefficient matrix on male trials in control group (median over sessions). MALE cells were detected from this matrix using change point analysis (80^th^ percentile, Figure S7A). OTHER cells were defined as all remaining cells. (**E**) Comparison of within-group correlation coefficients between MALE and OTHER cells in control sessions (each line is one session). Coefficients were Fisher-transformed for statistical testing. MALE cells showed more strongly correlated activity compared to OTHER cells (LME with MALE - OTHER coefficients as outcome variable: *Estimate* = 0.117, *SE* = 0.033, *t*(6.89) = 3.52, *P* = 0.00997**). (**F-G**) Same as (**D-E**) but now in Halo sessions. MALE and OTHER cells did not differ in their strength of correlated activity (LME with MALE - OTHER coefficients as outcome variable: *Estimate* = 0.038, *SE* = 0.053, *t*(7.00) = 0.72, *P* = 0.497). (**H-I**) Activity of MALE and OTHER cells on male (left) and milk (right) trials in control (**H**) and Halo (**I**) sessions. (**J**) Activity of MALE (left) and OTHER (right) cells on male and milk trials in control sessions. MALE cells significantly and stably distinguished between male and milk trials starting from 0.6 s before the run start, whereas OTHER cells did not distinguish trial outcome at any timepoint (LME with male - milk activity as outcome variable, and timepoint and cell group (MALE, OTHER) as fixed effects; significant timepoint x cell group interaction (*F*(15, 1085) = 49.14, *P* < 0.001***), followed by post hoc tests (difference from 0) at each timepoint in each cell group; black bar indicates *P* < 0.05 after Holm correction for multiple comparisons). (**K**) Same as (**J**), but now in Halo sessions. MALE cells briefly distinguished between male and milk trials at 0.6 s before the run start but only became stably significant starting from 0.4 s into the run. OTHER cells did not distinguish trial outcome at any timepoint (LME with male - milk activity as outcome variable, and timepoint and cell group (MALE, OTHER) as fixed effects; significant timepoint x cell group interaction (*F*(15, 1178) = 15.19, *P* < 0.001***), followed by post hoc tests (difference from 0) at each timepoint in each cell group; star and black bar indicates *P* < 0.05 after Holm correction for multiple comparisons). Throughout the figure, boxplots and thick lines/shaded areas indicate median and IQR over sessions.

If there were a dedicated ensemble of male-representing cells, we would expect them to prefer the male at the goal timepoint (as measured by the separability score) and show similar activity trajectories to each other on male trials. We therefore sorted cells by their separability values in each session. We then binned them according to percentiles (Figures 3B-C) and considered each bin’s activity trajectory over timepoints on male trials. We then correlated the activity trajectories between all bin pairs (Figure 3C) and visualized the correlation coefficients as a heat map (Figure 3D).

In the control group, we observed a strong boost in pairwise correlations at the 80^th^ percentile (Figures 3D-E), as detected using change point analysis (Figure S7A). This threshold corresponded to a positive (male-preferring) separability value of approximately 0.67 (Figure S7B; median over control sessions). We therefore defined MALE cells in each session as having a separability value at or above the 80^th^ percentile. We defined all remaining cells (below 80^th^ percentile) as OTHER cells. These results suggest that there is a dedicated ensemble of male-representing cells, which prefer the male at the goal timepoint and show correlated activity on male trials. These cells can be defined based on their ability to discriminate between male and milk trials at the goal timepoint.

To compare MALE and OTHER cells between control and Halo groups, we defined these cell types in Halo sessions using the same criteria (80^th^ percentile threshold on separability at goal timepoint, Figures 3F-G). Applying the same criteria to control and Halo sessions was justified because both groups had no manipulation of OXTR neurons at the goal timepoint (optogenetic light off; Figures 2J, S6B). Further, separability at the 80^th^ percentile was similar between control and Halo groups (Figure S7B).

Interestingly, in the Halo group, MALE and OTHER cells showed a similar level of correlated activity (Figures 3F-G). Indeed, activity appeared to be strongly correlated overall (Figures 3F-G). Widespread correlated activity was likely due to the optogenetic-induced increase in activity at pre-goal timepoints (Figures 2K-L), which may have dampened variability in activity trajectories over cells by pushing all activity towards a ceiling.

For completeness, we also explored activity correlations on milk trials (Figures S7C-F) and found similar effects to male trials. In the control group, MALE cells were more strongly correlated than OTHER cells (Figures S7C-D). In the Halo group, there was no difference between MALE and OTHER cells in their activity correlations (Figures S7E-F). Finally, unlike MALE cells, there was no obvious milk-representing ensemble in the control group, which would have appeared as a cluster of strongly correlated cells at the top left of the correlation matrix (Figure S7C) detected using change point analysis (Figure S7G). These results suggest that MALE and OTHER cells show similar correlation properties across trial types, and that the pyramidal population can best be described by two cell groups (MALE and OTHER).

### Inhibiting OXTR neurons disrupts the ability of MALE cells to represent the male option before the run

Now that we defined MALE and OTHER cells in each experimental group, we could track their activity over trial timepoints (Figures 3H-I). As expected, compared to OTHER cells, MALE cells in the control group ramped up their activity leading to the male goal on male trials (Figure 3H, left), but not to the milk goal on milk trials (Figure 3H, right). In the Halo group, ramping of MALE cells on male trials was less obvious because OTHER cells also showed elevated activity prior to the goal (Figure 3I).

To determine when MALE and OTHER cells began to distinguish trial outcome, we compared each cell group’s activity between male and milk trials at each timepoint (Figures 3J-K). In the control group, MALE cells showed significantly higher activity on male trials (compared to milk) starting at 0.6 seconds before the run start (Figure 3J, left). Even though OTHER cells slightly favored the milk outcome, this difference was not significant at any timepoint (Figure 3J, right). These results indicate that MALE cells reliably and preferentially represent the male option starting before the run.

In the Halo group, MALE cells briefly showed significantly elevated activity on male trials at 0.6 s before the run, before becoming non-significant at subsequent pre-run timepoints (Figure 3K, left). MALE cells resumed significantly elevated activity on male trials starting at 0.4 s after the run start. OTHER cells did not distinguish between trial types at any timepoint (Figure 3K, right). These results indicate that inhibiting OXTR neurons delays the appearance of a stable MALE representation from before to after the run start.

Together, these results suggest that MALE cells normally represent the male option during decision-making and that OXTR neurons stabilize this representation. This raises the possibility that OXTR neurons interact with MALE cells to promote male choice. To explore a potential mechanism for this interaction, we developed a computational model of decision-making.

### Asymmetric competition model reproduces experimental results

In a commonly used model of cortical decision-making, two excitatory cell groups, each representing a behavioral option, compete with each other in the presence of common inhibition to determine the behavioral outcome^37,38^. We therefore considered a competition between two pyramidal groups representing the male and milk options (Figure 4A). Both cell groups received inhibitory input from OXTR neurons, and featured recurrent, within-group connections among cells.

**Figure 4.**
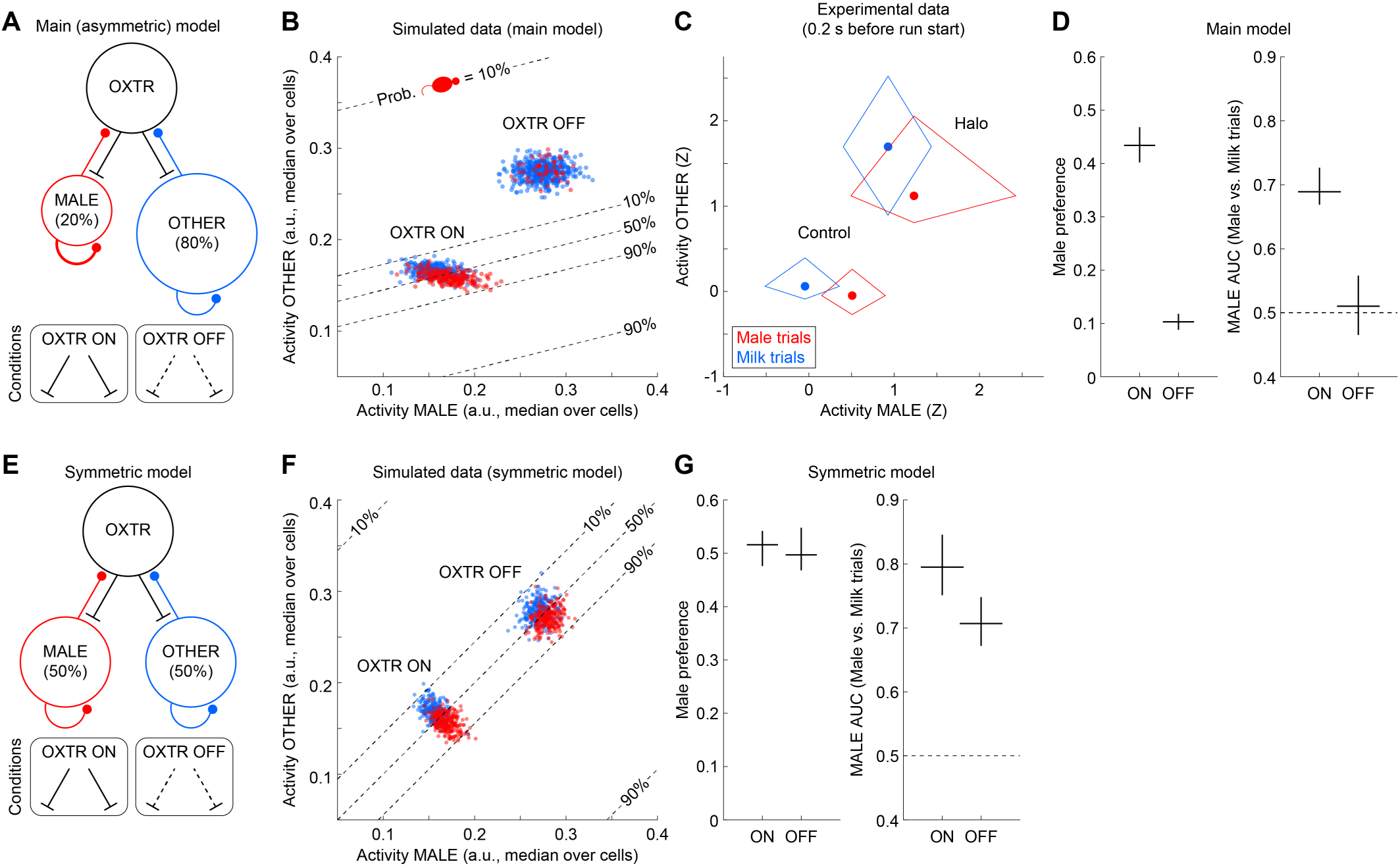
Computational model reproduces effects of inhibiting mPFC OXTR neurons on disrupting male choice and MALE cells’ representation before the run. (**A**) Model architecture, consisting of three cell groups: OXTR neurons, MALE cells, and all other (non-MALE) pyramidal neurons (OTHER cells). OXTR neurons inhibit MALE and OTHER cells, whereas MALE and OTHER cells excite OXTR neurons. MALE and OTHER cells also excite themselves through within-group connectivity, with MALE cells having stronger connectivity (thicker line). To reproduce experimental control and Halo conditions, the model was simulated with normal or reduced input from OXTR neurons (OXTR ON or OFF, respectively). (**B**) Representative simulation run of MALE and OTHER cells’ activity (*n* = 500 OXTR ON trials, 500 OXTR OFF trials). Outcome of each trial (male (red) or milk (blue)) determined by probability contours. (**C**) Experimentally recorded activity of MALE and OTHER cells on male and milk trials, plotted separately for control (*n* = 36) and Halo (*n* = 39) sessions. In each session, the activity of each cell group was summarized over male and milk trials separately, giving two values per session. The distribution of activity values over sessions for each trial type was plotted as a dot and boundary box (median and IQR over sessions). (**D**) Male preference (left) and AUC of MALE cells (right) on simulated OXTR ON and OXTR OFF trials. Horizontal and vertical lines show median and full range over *n* = 12 simulation runs. (**E**) Symmetric model architecture containing equally-sized MALE and OTHER cell groups. (**F-G**) Same as (**B, D**) but now using symmetric model.

When defining the two pyramidal groups, we split the pyramidal population to match the proportion of MALE cells observed in our experimental control data (Figure 3D). This gave a group of MALE cells (20%) representing the male option and a group of all non-MALE (OTHER, 80%) cells representing the milk option. Because we did not experimentally observe a dedicated milk-preferring ensemble (Figures S7C, G), we considered OTHER cells as representing milk as a “default choice.” This interpretation would not require a dedicated ensemble but would still be valid as a source of competition to the (non-default) male option.

We implemented behavioral choice in the model using a voting process, in which each cell (MALE or OTHER) contributed its activity as a vote for the male or milk option, respectively (see Methods). The difference in total (summed) activity between groups (MALE versus OTHER cells) could then be read out to make a behavioral choice. Because there were much fewer MALE cells than OTHER cells, MALE cells required a higher activity range to effectively compete with OTHER cells. We therefore specified stronger within-group connectivity of MALE cells (thick line self-connecting MALE cell group in Figure 4A), which allowed them to amplify their activity. Stronger MALE connectivity was consistent with our experimental observation that MALE cells’ activity was more strongly correlated compared to OTHER cells (Figures 3E, S7D).

We next simulated a network model with the proposed connectivity (Figure 4A) on individual trials (dots in Figure 4B). To reproduce our experimental conditions (control or Halo), we simulated trials with normal (“OXTR ON”) or reduced (“OXTR OFF”) input from OXTR neurons, effectively enabling or inhibiting OXTR neurons. Inhibiting OXTR neurons in the model increased pyramidal activity (Figure 4B), which qualitatively matched our experimental data before the run (Figure 4C). This indicated that the model reasonably captured OXTR neurons’ regulation of pyramidal activity at a putative decision-making timepoint.

We then determined each simulated trial’s outcome in a probabilistic manner based on the difference in total activity between MALE and OTHER cells (see Methods). The probability of male trials, set to range from 0.1 (low) to 0.9 (high), could be visualized as contours over the space of MALE and OTHER cells’ activity (Figure 4B). The slope of these contours reflected the relative proportions of MALE and OTHER cells and was shallow based on the much higher proportion of OTHER cells. This shallow slope indicated that male trials were most likely at low activity levels of OTHER cells. In that sense, OTHER cells gated the ability of MALE cells to drive male choice.

We then asked whether the model reproduced our experimental findings that inhibiting OXTR neurons disrupts male choice (Figure 1G) and MALE cells’ representation of the male option at a putative decision-making timepoint (before the run; Figures 3J-K). We applied the same metrics on simulated trials as we did for our experimental data: male preference (proportion of male trials) and the difference in activity between male and milk trials (summarized here as an ROC AUC value; Figure 4D). We performed multiple simulation runs to assess the consistency of these measures. Consistently over simulation runs, inhibiting OXTR neurons reduced both male preference and MALE AUC (Figure 4D), reproducing our experimental results.

Both of these results were due to increased activity of OTHER cells. Specifically, OXTR neurons normally regulated OTHER cells, such that variation in MALE cells’ activity could influence trial outcome (see OXTR ON trials in Figure 4B; x-axis shows variation in MALE cells’ activity over trials). This produced more male trials (higher male preference). It also dictated that MALE cells’ activity mapped onto outcome, leading to a greater discriminability between male and milk trials (higher AUC). On the other hand, inhibiting OXTR neurons increased OTHER cells’ activity, such that OTHER cells blocked the ability of MALE cells to influence trial outcome. This led to lower male preference and MALE AUC.

To compare our main model to a traditional models with equally-sized pyramidal groups^37,38^, we made a second model (“symmetric model”) with equally-sized MALE and OTHER groups (and equal connectivity within groups; Figure 4E). In the same fashion as our main model, we simulated OXTR ON and OXTR OFF trials, confirming that inhibiting OXTR neurons (OXTR OFF) increased overall activity (Figure 4F). We also tested how inhibiting OXTR neurons affected male preference and MALE AUC (Figure 4G). In contrast to the main model, inhibiting OXTR neurons did not obviously affect male preference. This was due to a balanced competition, in which increasing overall activity did not bias the competition towards either group of MALE or OTHER cells.

As in the main model, inhibiting OXTR neurons in the symmetric model reduced MALE AUC. However, the main underlying cause was different. While reduced AUC in the main model was primarily explained by the outsized influence of OTHER cells, reduced AUC in the symmetric model was due to a change in the shape of the activity distribution dictated by the model architecture. Normally, input from OXTR neurons regulated the relative activity of MALE and OTHER cells, such that the activity distribution took a line-like shape (see OXTR ON trials in Figure 4F). Inhibiting OXTR neurons pushed both MALE and OTHER cells’ activity towards their maximum values, producing a more circular shape (see OXTR OFF trials in Figure 4F). The line-like shape of OXTR ON trials better aligned to the activity axis (x-axis) of MALE cells, leading to a higher AUC value than on OXTR OFF trials. Given that we observed strong reductions in both male preference and MALE discriminability in our experimental data, the main model (Figure 4A) better matches our experimental data.

In summary, our model and data suggest that mPFC OXTR neurons normally regulate pyramidal population activity to stay within a specific range. Within this range, OTHER cells’ activity is low enough for variation in MALE cells’ activity to effectively drive male choice.

## Discussion

In this study, we report that mPFC OXTR neurons play a key role in choosing interaction with a male over an appetitive non-social alternative (milk) in female mice. Inhibiting OXTR neurons dampened male choice (Figures 1E-H), most strongly during sexual receptivity (Figure 1H). Inhibiting OXTR neurons also increased pyramidal population activity (Figure 2), which disrupted the ability of a small, specific cell group (MALE cells) to represent the male option during decision-making (Figures 3J-K).

Our computational model (Figure 4A) provided a potential mechanism by which OXTR neurons support male choice. Specifically, MALE cells compete against a much larger, but more weakly-connected, cell group (OTHER cells) representing the milk alternative. The larger group size of OTHER cells allows them to gate the competition: depending on their activity level, OTHER cells either permit (low activity) or preclude (high activity) MALE cells from competing. Common inhibition from OXTR neurons normally bounds both MALE and OTHER cells’ activity. Critically, this keeps OTHER cells in a permissive regime (low activity), where MALE cells’ activity can be sufficiently high to win the competition. In the absence of inhibition from OXTR neurons, elevated activity of OTHER cells precludes MALE cells from meaningfully driving male choice.

This model is consistent with previous literature on female rodent sociosexual behavior, showing that sociosexual motivation and behavioral expression are gated by multiple factors such as ovarian hormones, male odors, and courtship behaviors^39–41^. Our results suggest that a competition mechanism in mPFC gates females’ selection of a male over an appetitive non-social alternative. To what extent hormones such as estradiol interact with OXTR neurons and/or the pyramidal population to calibrate this competition is an exciting open question for future work. Further, given that oxytocin increases OXTR neurons’ activity (Figure S3B and previous work^22^) and is involved in sociosexual motivation and behavior^22,42,43^, it would be valuable to test whether modulating oxytocin action affects this competition and the ability of MALE cells to select the male option. This would contribute to understanding oxytocin’s role in sociosexual behavior, as well as more generally in behavioral flexibility^44^.

This competition framework implies that one or more downstream areas reads out the relative activity of MALE and OTHER cells from the mPFC and converts this difference into a behavioral choice. The mPFC projects to many brain areas including the amygdala, ventral tegmental area, and striatum^45^. Optogenetically activating OXTR interneurons in the infralimbic cortex has been shown to inhibit pyramidal neurons projecting to the basolateral amygdala (BLA) in rats^24^, suggesting that mPFC OXTR neurons regulate the mPFC-to-BLA pyramidal projection. Given the importance of reciprocal prefrontal-amygdala projections in social decision-making^8,46,47^, BLA may help to read out mPFC’s “vote” for the male or milk option in this task.

Our model also aligns with previous work suggesting that normal social functioning requires a balance of prefrontal excitation and inhibition^18–20^. Indeed, social deficits in autism and other disorders have been attributed to dysregulated prefrontal pyramidal activity^33,34^, including over-excited pyramidal activity^33^. One important consequence of dysregulated pyramidal activity could be that it alters the competition between subpopulations representing behavioral options. Indeed, our model shows that social behavior is at risk when it is represented by a small ensemble that competes against a much larger “default” pyramidal subpopulation.

Interestingly, autism features highly restricted and repetitive behaviors (e.g., body rocking, self-grooming)^48,49^. To what extent these can be considered “default behaviors” is an open question. However, the severity of these behaviors has been linked to reduced behavioral flexibility, suggesting that autistic individuals have difficulty switching from a preferred behavioral repertoire^50^. Future work could test whether reducing excessive pyramidal activity in autism models (e.g., *Cntnap2*-knockout^33^) unveils an ensemble representation of a social stimulus that predicts increased social behavior. Such a finding would be consistent with our competition model.

In conclusion, we developed a novel task of sociosexual decision-making in female mice and found that mPFC OXTR neurons play a key role in the behavioral expression and pyramidal representation of male choice. By modeling these results, we illustrated how OXTR neurons could promote male choice through a pyramidal competition mechanism. In addition to further characterizing this circuit in sociosexual behavior, future work should test whether such a competition mechanism applies to other interneuron types, social behavioral contexts, and disease contexts. Specifically, if dysregulating excitation/inhibition in mPFC biases pyramidal computation against small, highly-tuned ensembles, we would expect it to disrupt any behavior that requires such a dedicated ensemble. It could be that many social behaviors are at risk because they are represented this way.

## Methods

All animal procedures and experiments were performed in accordance with the guidelines of the Federal Veterinary Office of Switzerland and approved by the Cantonal Veterinary Office in Zurich, Switzerland.

### Mice overview

Subjects were female, heterozygous OXTR-Cre mice (Oxtr^tm1(cre/GFP)Rpa^/J, Jax: 030543), bred in our own facilities. They were socially-housed with at least one other female in individually-ventilated cages. The housing room had a 12-hour light/12-hour dark cycle (lights on from 6:30 AM to 6:30 PM) and was maintained at 21–24°C with 35–70% humidity.

### Experiments overview

Two main types of experiments were performed: 1) Population calcium imaging and optogenetics during sociosexual decision-making and 2) patch clamp electrophysiology and optogenetics. Methods associated with each experiment are described separately, as follows.

### Population calcium imaging and optogenetics during sociosexual decision-making

#### Mice

All subjects were confirmed by genotyping (Protocol 31289: Separated PCR Assay; The Jackson Laboratory) to be heterozygous Cre-expressing (Cre/+). The only exception was one control subject, which was Cre-negative. This subject did not express the Cre-dependent control fluorophore in OXTR neurons (see “Viral injection” section below) and only expressed GCaMP6m in pyramidal neurons. This subject was included in the control group because, like all other control subjects, it received the same experimental procedures (including viral injections) and would not be expected to show an effect of optogenetic light. In total, there were 8 control and 8 Halo subjects. Subjects were between 97 and 130 days old at the start of experiments (first handling session) and had a maximum age of 152 days by the end of experiments.

Social stimuli were C57BL/6J adult males and adolescent females. These animals were ordered from Charles River Laboratory Germany or Janvier Labs and group housed in our facilities. Upon arriving at our facilities, they were allowed at least 1 week to habituate before starting experiments. Males were between 63 and 112 days old during experiments, and females were between 25 and 38 days old. In using different ages for the two stimuli types, we aimed to match traditional social paradigms involving male subjects, in which the subject is exposed to an adult opposite-sex conspecific as a sociosexual stimulus^51,52^ and a young same-sex conspecific (∼3-6 weeks of age) as a non-sexual social stimulus^53–55^.

#### Experimental setup

The setup consisted of a custom Y-shaped arena (Maze Engineers), with three rectangular arms (35 cm length x 7 cm width x 20 cm height) and a triangular center chamber (7 cm edges). Two of the arms extended onto a platform, upon which the social stimulus chamber or milk dispenser could be interchangeably placed (“social and milk arms”). These arms also had distinct ground textures to help the subject differentiate between them. The third arm (“start arm”) had a speaker built into the wall, which interfaced with a function generator (Hantek HDG2022B) to produce a tone at the start of each trial. An air pressure system controlled the movement of five doors individually, three at the center of the arena (connecting the center triangle to each arm) and two at the end of the social and milk arms (in front of the social chamber and milk dispenser). The doors were elevated to the top of the arena at rest but could be triggered to lower down to the floor (open), allowing subjects to pass through the center chamber or access to the social stimulus/milk dispenser. An infrared sensor was placed at the landing position of one of the center doors, such that the exact timing of doors opening could be detected on each trial (beam break).

The milk dispenser consisted of a 50 mL syringe chamber (Braun Omnifix Luer Lock Solo) connected with tubing to a solenoid valve (SV74; Valcor Scientific) and blunt syringe needle (Z261378; Sigma-Aldrich). The dispenser was vertically clamped in place such that the needle threaded through a metal sipper into the arena at a mouse-accessible height (approximately 1 cm from floor). The milk solution was a 7.1% (by volume) dilution of condensed milk (Migros Kondensmilch gezuchert) in drinking water from the mouse housing facilities. Before each experiment, the solenoid opening time was calibrated to deliver approximately 8 mg of milk solution as a suspended droplet on a given trial.

The social chamber was a rectangular box (9 cm x 7 cm width x 20 cm height) with three closed sides and one open side with spaced bars (0.7 cm width with 0.9-1.3 cm spacing between bars). Immediately before the experiment, a novel male and female were removed from their home cages and placed in dedicated social chambers. Then, depending on the stimulus order in a given experiment, the male or female chamber was placed at the end of the social arm, with the open side facing the arm to allow for social (sniffing) interactions with the subject. The other chamber remained at a different location in the experimental room until it was swapped onto the arena half-way into the session.

The Inscopix nVoke system was used for calcium imaging and optogenetics. This system consisted of a miniaturized microscope (1.8 g), with two light channels for calcium imaging (455 +/- 8 nm, 0.5-1 mW/mm^2^) and optogenetics (620 +/- 30 nm, 10 mW/mm^2^). The microscope interfaced with a data acquisition (DAQ) box, which directly controlled the microscope, collected and stored imaging data (frames, 20 Hz), and outputted TTL synchronization pulses corresponding to these frames. To prevent the microscope cable from becoming tangled during experiments, the DAQ was placed on a custom-built rotating platform above the arena. Because the DAQ was powered by a portable power bank (xtorm AC Power Bank Pro), it could be rotated to untangle the cable.

ANY-maze video tracking software (version 6.06) and custom Arduino and Windows executable scripts were used to run experimental protocols and create all experimental files except for imaging movies (Inscopix DAQ). The software interfaced with a video camera (DMK 22BUC03 USB CMOS camera with T3Z 3510 CS vari-focal lens; The Imaging Source) above the arena for tracking animals’ center of mass and a custom air pressure control box (Maze Engineers) for lowering and elevating arena doors. It also interfaced with an A-Mi2 Digital Interface box (ANY-maze) for triggering the Inscopix DAQ (initiating neural recording and optogenetic light delivery), function generator (producing a tone), and Arduino Uno microcontroller (opening solenoid for milk delivery). Through its connection with the Digital Interface box, ANY-maze also received the TTL synchronization pulses from the Inscopix DAQ and detection of the center doors lowering from the infrared sensor and Arduino Uno. ANY-maze then stored these events alongside positional tracking and behavioral videos, such that neural and behavioral data could later be aligned on a common (ANY-maze) clock.

*Surgical procedures*. 1) Anesthesia: For viral injection and guide tube implantation procedures, subjects were preemptively treated with buprenorphine (Bupaq, 0.1 mg/kg; Streuli Pharma AG) 20 to 30 minutes prior to anesthesia. Anesthesia was induced using a ketamine-xylazine cocktail (90 mg/kg ketamine (Ketanarkon) and 8 mg/kg xylazine; Streuli Pharma AG), and the subjects were positioned in a stereotactic frame (Kopf Instruments). During the procedure, subjects were supplied with 95% medical oxygen (PanGas, Conoxia) via a face mask, and their body temperature was maintained at 37 °C using a temperature controller and heating pad. Their eyes were protected using eye ointment (Hylo Night; Hylo).

2) Viral injection: Once subjects were fully anesthetized, they were placed in the stereotactic frame. Following hair removal (Veet), cleaning (Betadine solution) and incision of the scalp, a 0.5 mm diameter hole was drilled through the skull (19007-05 micro drill burr; FST) above the right mPFC (anterior-posterior (AP) from Bregma: 1.9 mm, medial-lateral (ML): 0.3 mm). An injection needle (33 gauge Flexifil beveled needle; World Precision Instruments) was connected to a syringe (10 µL Nanofil; World Precision Instruments) and lowered into the mPFC (dorsal-ventral (DV): 2.1-2.2 mm from skull surface). Adeno-associated viruses (AAVs; see below) were then injected intracranially into the mPFC (Micro 4 MicroSyringe Pump; World Precision Instruments). The total injection volume was 500 nL, injected at a rate of 100 nL per minute. At the end of injection, the needle was left in place for 5 minutes to allow the viruses to diffuse into the tissue. After removal of the needle, the incision was closed with sutures.

Control subjects were injected with a mixture of ssAAV5/2-mCaMKIIα-GCaMP6m-WPRE-SV40p(A) (250 nL, Titer: 3.15 × 10^12^ vg/mL; UZH Viral Vector Facility), expressing GCaMP6m in pyramidal neurons, and rAAV5-EF1α-DIO-mCherry-WPRE (250 nL, Titer: 2.65 × 10^12^ vg/mL; UNC Vector Core), expressing mCherry exclusively in Cre-positive neurons.

Halo subjects were injected with a mixture of ssAAV5/2-mCaMKIIα-GCaMP6m-WPRE-SV40p(A) (250 nL, Titer: 3.15 × 10^12^ vg/mL; UZH Viral Vector Facility), expressing GCaMP6m in pyramidal neurons, and rAAV5-EF1α-DIO-eNpHR3.0-mCherry-WPRE (250 nL, Titer: 1.65 × 10^12^ vg/mL; UNC Vector Core), expressing Halorhodopsin and mCherry exclusively in Cre-positive neurons.

3) Guide tube implantation: After 7 to 16 days of recovery from the viral injection surgery, subjects received a second surgery implanting a guide tube directly above the viral injection site. The guide tube consisted of a stainless steel cannula (1.2 mm diameter; Ziggy’s Tubes and Wires) with a custom glass coverslip (0.125 mm-thick BK7 glass; Electron Microscopy Sciences) glued to the end, as previously described^56^. A circular craniotomy with a diameter of 1.2 mm was performed above the mPFC (AP: 1.9 mm, ML: 0.3 mm). Then, tissue was aspirated to a depth of ∼1.9 mm from the skull surface, and the guide tube was lowered to a depth of 1.9 to 2.2 mm from the skull surface.

The guide tube was secured to the skull using UV-curable glue (4305 LC; Loctite). To further stabilize the implant, a layer of dental acrylic (Metabond; Parkell or Scotchbond ESPE; 3M) was applied. A custom metal bar was attached to the implant with dental acrylic cement (Paladur). This metal bar would later be used during experiments to secure the subject’s head when connecting the microscope.

4) Postsurgical analgesia: For four days including the surgery day, subjects received buprenorphine subcutaneously (0.1 mg/kg) every 6 to 8 hours during the light cycle and in the drinking water (0.01 mg/mL) during the dark cycle. Additionally, subjects received carprofen subcutaneously (Rimadyl, 4 mg/kg; Zoetis) every 12 hours.

5) Baseplate mounting: After 24 to 41 days of recovery from the guide tube implantation surgery, subjects were anesthetized and placed in the stereotactic frame. A gradient-index (GRIN) lens (1 mm diameter, 4.2 mm length, GT-IFRL-100-101027-50-NC; Grintech) was inserted in the guide tube and secured with glue (Flow-It; Pentron). A microscope baseplate was then secured above the lens using glue (Flow-It).

#### Experiment timeline

The decision-making task was the last phase in a seven-phase timeline for each subject, as shown in Figure S4A and described as follows.

In all phases, the home cages of the subject and any social stimuli were first transferred from the housing room to the experimental room. Animals were habituated to the room for at least 15 minutes (Handling, Microscope habituation, Drink training sessions; see below) or at least 30 minutes (Arena habituation, Arena training, Decision-making task sessions) in their home cages. At the end of each session, the subject and stimulus home cages were brought back to the housing room.

1. Handling (4-5 daily sessions): The subject was gently held by the experimenter for approximately 1-5 minutes, before being returned to its home cage.
2. Microscope habituation (3 daily sessions): The subject was transferred to the top of a running wheel, and its head was secured in place by clamping the metal bar on the implant. This setup allowed subjects to comfortably run on the wheel with a fixed head position for microscope mounting. The implant and exposed top of the lens was gently cleaned with a Q-tip. Then a plastic (“dummy”) microscope of similar size and weight to the actual microscope was mounted to the implant. The subject was released from the mounting station, transferred to an open, empty cage, and allowed to freely move with the dummy microscope for 5 minutes. Then, the subject was returned to the running wheel, and the dummy microscope was removed.
3. Measurement of baseline body weight: The subject’s body weight was measured on the third session of microscope habituation. This measurement was used as the baseline weight. The water bottle was removed from the cage on the same day (see *Water scheduling and weight tracking* below).
4. Drink training (2 daily sessions): The dummy microscope was mounted as described above. The subject was then transferred to a plastic cylindrical chamber (18.5 cm diameter, 29 cm height) at a separate location from the decision-making arena. On one side of the chamber, the tip of a milk dispenser (same design as in decision-making arena) entered into the chamber through a small hole. Subjects then had free access to lick milk from the dispenser for up to 15 minutes. Then the subject was returned to the running wheel, and the dummy microscope was removed.
5. Arena habituation (1 session): After the second session of drink training (same day, in the afternoon), the subject was brought back to the experimental room from the housing room. The real microscope was mounted for imaging, and the subject was transferred to the start arm of the decision-making arena, where it had 15 minutes to freely explore the three arms and center chamber. The social stimulus and milk dispenser were not present during this habituation session. Then the subject was returned to the running wheel, and the microscope was removed.
6. Arena training (3 daily sessions): The microscope was mounted as described above, and the subject was transferred into the start arm of the arena. The subject was then exposed to the task design (described in detail below; milk available from dispenser), with two differences. First, the subject encountered an empty social chamber at the end of the social arm (no social stimulus inside). Second, the session duration was bounded by time (within 1 hour 30 min.), rather than trials because subjects were not yet expected to stably perform the task.
7. Decision-making task (7-12 daily sessions): The microscope was mounted, and the subject was transferred to the start arm of the arena. The subject then completed up to 32 trials of the decision-making task. In early sessions before introducing optogenetic light, the experimenter could decrease the required number of trials if it was clear in the first ∼30 minutes that the subject was still learning the task and would not complete 32 trials in a reasonable timeframe (target of 1 hour). In this case, the experimenter planned a conservative and even number of trials (between 22 and 30) to equally expose the subject to the male and female stimuli (each for half the number of trials). The only exceptions were 2 early sessions, in which the subject was exposed to only one of the two stimuli due to experimental problems (especially slow performance of the subject or technical issues with the milk dispenser), which limited the possible trial count to 10 and 14 trials, respectively. Once subjects began optogenetic sessions, they completed 32 trials per session.

Each trial proceeded as follows: the subject began in the start arm, with the three center doors elevated (closed) to prevent entry into the center chamber. A tone played from the speaker on the start arm wall (5 s, 4 kHz, 90 dB(A)), and the three center doors lowered to the arena ground. This allowed the subject to enter the social or milk arm (free choice). Once the subject entered its chosen arm, the two center doors facing the start arm and non-chosen option elevated, preventing re-entry to the start arm or a switch in choice to the other option. Then, as the subject ran down the chosen arm towards the social or milk goal, the third center door connecting the chosen arm and center chamber elevated, preventing re-entry to the center chamber. Once the subject entered the goal area in front of the social stimulus or milk dispenser (7 cm x 8.5 cm region at end of social or milk arm), it had 15 s to socially interact or lick a milk droplet suspended from the tip of the milk dispenser. Then, a door in front of the social chamber or milk dispenser closed, preventing further interaction or licking. To allow the subject to return back to the start arm, two of the center doors opened (one connecting the chosen arm to the center chamber, the other connecting the center chamber to the start arm). Once the subject returned back to the start arm, these two center doors re-elevated, and the door in front of the chosen option re-lowered. This brought all doors back to their starting positions. If the subject had chosen the milk option, a new milk droplet was generated by triggering the solenoid valve in the milk dispenser. The inter-trial interval was randomly sampled within the range of 20 to 40 s.

#### Estrous cycle tracking

Each subject’s estrous cycle was tracked daily from microscope habituation onward. At the end of each session, a vaginal sample was collected as previously described^29^. Using a pipette, the vaginal opening was flushed with 0.015 mL double-distilled water (ddH_2_0). The sample was then placed onto a glass slide to air dry at room temperature. Once the sample fully dried, it was stained with a 0.1% crystal violet solution, coverslipped, and observed under a light microscope (Bresser LCD Micro 5MP). The estrous phase (proestrus, estrus, metestrus, diestrus) was evaluated based on the distribution of nucleated epithelial cells, cornified epithelial cells and leukocytes^29^. Given that mating in mice is more likely during proestrus (many nucleated epithelial cells) and the earlier part of estrus (some nucleated epithelial cells among cornified epithelial^28^)^31^, estrus was split into two phases (early, late) based on the presence or absence of nucleated epithelial cells. Early versus late estrus thus defined the distinction between putatively sexually receptive (proestrus, early estrus) and non-receptive sessions (all other phases). It is possible that sampling with ddH_2_0 (as opposed to saline) reduced the detectability of neutrophils due to cell rupturing^27^. However, this likely would not have affected the distinction of putatively sexually receptive sessions, which relied more strongly on nucleated and cornified epithelial cells.

#### Water scheduling and weight tracking

The water bottle was removed on the same day as the third session of microscope habituation. On each following day up to the last session of the decision-making task, the subject was given water access after the experimental session, once the subject had been returned to the housing room. A water bottle was placed in the subject’s home cage for at least 1 hour, allowing free drinking. Water access could be up to 2 hours on early-session days to allow subjects to habituate to the scheduling protocol. By the first day of the decision-making task, the daily water access was consistent at 1 hour.

To track each subject’s body weight over sessions, weight was measured at the start of each experimental session and compared to the baseline measurement, as follows:

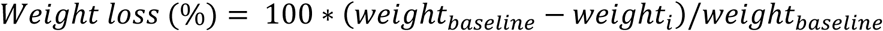

where *i* refers to a given session.

#### Histology

1. Perfusion: At the end of experiments, subjects were given a terminal dose of pentobarbital (Esconarkon, 200 mg/kg; Streuli Pharma AG). After loss of the pedal withdraw reflex, subjects were transcardially perfused with phosphate-buffered saline (PBS), followed by 4% paraformaldehyde (PFA). The implant was then removed, and the brain was post-fixed for 48 hours. It was then coronally sliced over the anterior/posterior range of the mPFC using a vibratome (VT1000 S; Leica) at a thickness of 50 μm. Slices were stored in PBS until further processing.
2. Verification of implant placement: Every third mPFC brain slice was Nissl stained (NeuroTrace 640/660; Invitrogen). Nissl staining highlights cytostructural differences within the tissue, helping to align the section to the atlas. The staining was performed following the protocol from Invitrogen, using a dilution of 1:50. After staining, the slices were mounted onto glass slides, and fluorescence images were taken using an Olympus BX51 microscope. The images were compared to the mouse brain atlas^57^, and the location of the implant base was marked for each subject (Figure S1F). Additionally, the presence of the mCherry tag in Cre-positive neurons was confirmed for each animal.
3. Immunofluorescence staining: Tissue slices were incubated overnight at 4°C with either a rabbit anti-Neurogranin primary antibody (AB5620; Merck Millipore; 1:2000) or a rabbit anti-GAD 65 & 67 primary antibody (AB1511; Merck Millipore; 1:500). They were subsequently incubated for 2 hours at room temperature with a secondary antibody, goat anti-rabbit Alexa Fluor 594 (A-11012; Invitrogen; 1:200). The slices were then stained with DAPI (D1306; Invitrogen; 1:1000) in 0.1 M PBS for 4 minutes before being mounted onto glass slides. Confocal images were captured (Leica Stellaris 5) using the red channel (594 nm; Neurogranin or GAD), the green channel (488 nm; GCaMP6m), and the blue channel (390 nm; DAPI). The images were viewed in the LAS-X software to evaluate overlap in labeling.

### Patch clamp electrophysiology and optogenetics

#### Mice

Subjects were between 89 and 138 days old on the day of experiments.

#### Viral injections

The same viral injection procedures were performed as described above (“Population calcium imaging and optogenetics during sociosexual decision-making”). Subjects were injected with a mixture of ssAAV5/2-mCaMKIIα-GCaMP6m-WPRE-SV40p(A) (250 nL, Titer: 3.15 × 10^12^ vg/mL; UZH Viral Vector Facility), expressing GCaMP6m in pyramidal neurons, and rAAV5-EF1α-DIO-eNpHR3.0-mCherry-WPRE (250 nL, Titer: 1.65 × 10^12^ vg/mL; UNC Vector Core), expressing Halorhodopsin and mCherry exclusively in Cre-positive neurons.

#### Brain dissection and acute slice preparation

5 to 6 weeks after viral injections, subjects were deeply anesthetized using isoflurane and rapidly decapitated. Brains were quickly removed and submerged in ice-cold artificial cerebrospinal fluid (aCSF; containing, in mM: 75 sucrose, 10 glucose, 80 NaCl, 26 NaHCO_3_, 7 MgSO_4_, 2.5 KCl, 1 NaH_2_PO_4_, 0.5 CaCl_2_) fumigated with a gas mixture of 95% O_2_ and 5% CO_2_. Coronal slices (300 μm for validation of OXTR neurons and 350 μm for paired recordings) were prepared from the mPFC (approximately 1.5 to 3 mm anterior to Bregma) using a vibratome (VT1200 S; Leica). Slices were incubated in oxygenated aCSF recording solution (containing, in mM: 15 glucose, 120 NaCl, 26 NaHCO_3_, 0.9 MgSO_4_, 2.5 KCl, 1.25 NaH_2_PO_4_, 1.3 CaCl_2_) at 36 °C for 15 minutes before being transferred to room temperature for at least 40 minutes prior to recordings.

#### Recording setup

During recordings, brain slices were held in oxygenated aCSF recording solution at 36 °C. Neurons were visualized using differential interference contrast (DIC) microscopy (Olympus BX61 WI) with a 60× water-immersion objective. Images were captured with a CellCam Kikker MT100 (Cairn). Patch pipettes with resistances of 4 to 6 MΩ were pulled from borosilicate glass and filled with intracellular solution (in mM: 10 HEPES, 20 KCl, 117 K-gluconate, 4 Mg-ATP, 0.3 GTP, 10 Na-phosphocreatine, pH adjusted to 7.3 with KOH). For morphological reconstruction, 0.2% biocytin was included in the pipette solution. Whole-cell somatic recordings were obtained in current-clamp or voltage-clamp mode using a MultiClamp 700A amplifier (Axon Instruments), digitized with a DigiData 1550B system, sampled at 50 kHz, and filtered at 3 kHz. Recordings were excluded if access resistance changed by more than 15% during the experiment. Bridge potentials were compensated, and liquid-junction potentials were not corrected. Optogenetic light was delivered to the sample using a Lumencor SPECTRA Light Engine (635 +/- 20 nm, 0.9-1.3 mW/mm^2^).

#### Validation of OXTR neurons

Halorhodopsin-expressing OXTR neurons (*n* = 18) were identified by mCherry fluorescence using a Lumencor SPECTRA Light Engine at 575 +/- 25 nm. Resting membrane potentials ranged from -65 mV to -80 mV, and access resistances were between 15 and 25 MΩ. After establishing a stable membrane potential, depolarizing and hyperpolarizing current steps were applied to characterize neuronal properties, including spike patterns and rebound undershoots during hyperpolarization. Optogenetic light (5 repetitions of 1.5 s duration) was delivered to confirm functional Halorhodopsin expression (hyperpolarization response).

Next, in 12 of 18 neurons (“aCSF + OXT cells”), oxytocin (OXT) was added to the aCSF solution during membrane potential recordings. OXT was added by exchanging the original aCSF perfusate with a second aCSF solution containing 100 nM OXT (O3251; Sigma-Aldrich; concentration matching previous work^22^). The remaining 6 neurons only received aCSF (“aCSF only cells”), but were experimentally matched to aCSF + OXT cells by exchanging the original perfusate with a second identical solution (no OXT). Exchanging the perfusate produced an air bubble leading to the recording chamber, which was used to define a wash-in start time (bubble entering recording chamber). Neuronal responses were recorded for at least 12 minutes from this timepoint.

To quantify cells’ responses to OXT, two 180 s-long windows were defined immediately before the start of wash-in (“baseline window”) and in the last part of wash-in (“active window”) for each cell. The membrane potential was low-pass filtered (Butterworth 2^nd^ order filter with 2 Hz cutoff; *butter* function in SciPy (version 1.11.4) library in Python (version 3.1)) and averaged over samples within each window, producing one baseline and one active value. The change in membrane potential (active - baseline windows) was computed for each cell.

#### Paired recordings of OXTR neurons and pyramidal neurons (PNs)

As described above, Halorhodopsin-expressing OXTR neurons were identified (mCherry fluorescence), patched, and validated (hyperpolarization response to optogenetic light). Based on previous work showing monosynaptic connections of SST interneurons onto layer 2/3 (L2/3) PNs in mouse frontal cortex^32^, OXTR neurons next to viable L2/3 PNs were targeted. Upon patching an OXTR neuron, a second patch was established in an adjacent L2/3 PN, confirmed by its morphology and spiking patterns. Whole-cell recordings were then obtained from both cells. To assess monosynaptic connectivity in a consistent way to previous work^32^, the PN was voltage-clamped at +40 mV, while the OXTR neuron was stimulated to fire 3 to 5 high-frequency action potentials via current injection. In 3 out of 11 neuron pairs (27.3%), postsynaptic current responses were observed in the PN. These pairs were thus used for further recording in current clamp, as follows.

In a given pair, a current injection level to each of the PN and OXTR neurons was calibrated, such that the PN produced multiple but few action potentials just above firing threshold, while the OXTR neuron produced multiple high-frequency action potentials. Then, a stimulation protocol was performed, consisting of three 6 s-long phases spaced by 0.5 s: (1) PN current injection, (2) both PN and OXTR neuron current injection, and (3) both PN and OXTR neuron current injection combined with optogenetic light delivery. The number of action potentials of each cell was counted in each phase. This protocol was performed twice in a row in two cell pairs, and four times in the third cell pair (repetitions). Action potential counts were averaged over repetitions, giving three final values per cell over the three phases.

#### Histology

After recordings, cells were gently de-patched, and slices were allowed to rest for 10 minutes before being fixed in 4% PFA in PBS for 24 hours. Biocytin-streptavidin staining^58^ was performed to fluorescently label and reconstruct the recorded PN and OXTR neuron pair. Stained sections were imaged using a Leica Stellaris 5 confocal microscope and overlaid on the mouse brain atlas^57^. The location of the recorded cell pair was mapped to the mPFC (prelimbic or cingulate cortex).

### Analyses (sociosexual decision-making task)

Behavioral and neural data analyses were performed in Matlab (versions R2018B and R2025B), unless otherwise noted.

#### Data selection

Analyses were performed on decision-making task sessions (last phase in experiment timeline, described above). Five sessions were excluded from all analyses due to technical issues with optogenetic light delivery. An additional 2 sessions had technical problems with imaging (occluded or unstable field of view) and thus were excluded from all analyses except for return latency (Figures S4E-G). In the analysis fitting a baseline (pre-opto.) curve onto female versus male preference values (Figures 1E, S5A-B), 2 sessions were additionally excluded due to especially slow performance of the subject or technical issues with the milk dispenser, which severely limited the number of available trials (subject was exposed to only one of the two social stimuli).

Within each session, all available trials were used for behavioral and neural analyses. The only exceptions were 7 trials (from a total of 6 sessions), which were excluded due to the subject not reaching the social stimulus or milk dispenser, the subject not drinking from the milk spout, or disrupted tracking/control of the maze doors.

#### Behavior

The following behavioral analyses were performed.

1. Male and female preference (each session). Trials from each session were categorized by their outcome as social or milk. Given that each session included both a male and female stimulus (each for half of the total number of trials, typically 16 out of 32), trials were further categorized as male, milk in the presence of the male, female, and milk in the presence of the female. Male preference was computed as the proportion of male trials, considering male trials and milk trials in the presence of the male. Female preference was computed as the proportion of female trials, considering female trials and milk trials in the presence of the female. Preference ranged from 0 to 1, where 0 indicated all milk trials, 0.5 indicated equal numbers of social and milk trials, and 1 indicated all social trials (“social” here refers to male or female, depending on whether male or female preference is calculated).
2. Positional tracking and preprocessing (each session). Each subject’s center-of-mass position (XY pixel coordinates) was tracked over each session (13.91 to 15.03 Hz) and interpolated to match the neural data (5 Hz; see *Calcium signal preprocessing* below). Position was then projected onto an axis connecting the social and milk goal locations, specifically the midline entry points into each goal area (see Figure 1I). The goal axis ranged from -1 to 1, where -1 corresponded to the milk goal, 0 to the arena midline, and 1 to the social goal. As subjects passed through the entry points and reached the social chamber or milk dispenser, the projected position was capped at 1 or -1. All subsequent position-based analyses (velocity, alignment point, run definition) used the projected position. The only exception was movement speed during optogenetic light (see below), which used the position in XY coordinates.
3. Velocity (each session). Using the projected position over time samples in each session, velocity was calculated at each time sample as the change in position from the current to next time sample, divided by the time interval (0.2 s).
4. Alignment point and run start (each trial). On each trial, the time at which the subject entered the social or milk arm was detected based on center-of-mass tracking (ANY-maze). Projected position samples were then taken within a window of 2.8 s before to 0.3 s after arm entry. Small, non-directed fluctuations in the position were smoothed by taking a 0.6 s moving average within this window. Working backward in time from arm entry, the first sample at which the subject’s position monotonically increased toward the social goal or decreased toward the milk goal was defined as the alignment point. If a monotonic change in position only occurred once the subject entered the social or milk arm, the first time sample at or directly after arm entry was used as the alignment point. The run start was defined as 0.6 s before the alignment point.
5. Optogenetic light duration (each trial and session). Optogenetic light was triggered on each trial from the second half of the tone to entry into the social or milk arm, as detected and controlled by the ANY-maze tracking system. The duration of optogenetic light was computed from the ANY-maze experimental record. Light duration was summarized over trials in each session (median) to obtain one value per session.
6. Movement speed during optogenetic light (each trial and session). Position in XY pixel coordinates was extracted during the first 2.5 s of optogenetic light, giving a series of position samples spaced in time (5 Hz). The Euclidean distance between samples was summed and converted from pixels to cm based on the ANY-maze calibration setting (0.15 cm per pixel). To obtain speed, the total distance travelled was divided by the corresponding time duration (sum of time intervals between samples). Speed was summarized over trials in each session (median) to obtain one value per session.
7. Trial timepoints (each trial). On each trial, specific timepoints were defined for neural and behavioral analyses. These timepoints consisted of adjacent time samples around the run start (from 0.6 before to 1.8 s after the run start) as well as 2.5 s-long windows containing multiple samples. These windows were immediately before the tone start, the first half of the tone before optogenetic light delivery, the second half of the tone with optogenetic light on, and entry into the goal area (“goal”). While some timepoints were defined to be distinct (e.g., adjacent timepoints around the run start), others could at least partially overlap (e.g., late run timepoints and goal). To determine how often timepoints overlapped, timepoints were compared in a pairwise fashion (e.g., 1.8 s after run start vs. goal) on each trial. For each pair, the proportion of trials in which the timepoints at least partially overlapped was computed. The timing of optogenetic light was also compared to each timepoint over trials. The proportion of trials in which the optogenetic light at least partially overlapped with each timepoint was computed.
8. Return latency (each trial and session). The return latency was computed on each trial based on ANY-maze tracking of the subject and arena doors. It spanned from the center doors fully lowering at the end of the trial (detected by an infrared beam break sensor) to the subject returning to the start arm (see Figure S4E). Latencies were summarized over trials in each session (median) to obtain one value per session.

#### Neural data preprocessing

Calcium imaging data was preprocessed using the Inscopix Data Processing software (IDPS, version 1.8). For each subject, calcium imaging movies (20 Hz) from all sessions were imported. For each session, the movie was spatially cropped at its edges to remove dark or occluded non-fluorescing parts of the field of view. It was then spatially and temporally downsampled, each by a factor of 4. This reduced the frame rate to 5 Hz. The movie was then spatially filtered (spatial band-pass filter with 0.005 low and 0.5 high pixel^-1^ cutoffs) and motion-corrected relative to the first frame (maximum translation of 20 pixels). The ΔF/F trace of each pixel was computed over all frames using the mean frame as a baseline. The PCA-ICA method was used to identify candidate cells. The ΔF/F trace of each candidate cell was extracted using the *Apply Contours* function.

Once candidate cells were identified in each session, they were longitudinally registered over all sessions. Each candidate cell was then manually checked. A candidate cell was accepted if it showed a typical activity profile of the GCaMP6m indicator (peaks with fast rise and slow decay), a round cell image, and minimal activity interference from adjacent cells.

Cells’ ΔF/F signals in each session were then imported into Matlab, where they were referenced to the ANY-maze clock using the common TTL synchronization pulses. Neural data was then parsed into trials using the ANY-maze behavioral records. On each trial, each cell’s ΔF/F signal was Z-scored relative to a window at the trial start (2.5 s window before tone start, mean and standard deviation of all samples within window; see Figure 2J).

#### Cell maps

Cell images were exported from IDPS and stacked in ImageJ (version 1.54f). The stack was then maximum-intensity projected to a single image.

#### Single-cell and population analyses

The following analyses were performed on Z-scored neural activity.

1. Single-cell responses to optogenetic light (each session). On each trial of a given session, windows of 2.5 s duration were defined immediately before and at the start of optogenetic light (“Pre-light” and “Light start”, respectively). Then, the activity of each cell was taken within the two windows. This gave multiple time samples for a given cell within each window, over which the activity was averaged to obtain one Pre-light and one Light start value on each trial. The change in activity from Pre-Light to Light start was evaluated over trials for each cell using a Wilcoxon signed-rank test. Cells’ *P* values were then corrected for multiple comparisons using the Benjamini-Hochberg method (*fdr_bh* function^59^ in Matlab; false discovery rate: 0.05). Cells that showed a significant increase or decrease in activity (*P* < 0.05) were labeled as such and counted.
2. Population response to optogenetic light (each trial and session). The activity of each cell was extracted in the Pre-light and Light start windows on each trial, as described above. Activity in each window as well as the change in activity from Pre-light to Light start were summarized over cells (median) on each trial. These values were then summarized over trials (median) in each session to obtain one value (per window) per session. In addition to the Pre-light and Light start windows (defined based on light start), the same analyses were also performed on two 2.5 s-long windows immediately before and after the light turned off on each trial.
3. Activity of individual cells over trial timepoints (each trial and session). Trial timepoints were either individual time samples or windows containing multiple samples (see definition above). In the case of individual time samples, the cell’s activity was indexed at each sample. In the case of windows, the cell’s activity was averaged over samples in each window, giving one value per window. This gave a 1 x timepoints vector of activity values for each cell on each trial. When plotting the activity of individual cells over timepoints on a specific trial type (e.g., male trials), the activity of each cell was summarized (median) over trials of that type in each session. When plotting the activity of the population or specific cell group over timepoints on a specific trial type, activity was summarized over cells (median) on each trial and then summarized over trials (median) in each session.
4. Separability score (all individual cells in each session). For each cell in a given session, its activity was taken at the goal timepoint on male and milk trials. This gave two sets of values (male versus milk trials), from which the ROC AUC (0 to 1) was computed. The AUC was then rescaled between -1 to 1 (“separability”; 2 x (AUC - 0.5)). A separability value of 1 indicated maximal activity preference for the male (activity on male trials above all milk trials), and a value of -1 indicated maximal activity preference for the milk (activity on milk trials above all male trials). Within each session, cells were sorted by their separability values, and the percentiles of the separability distribution for that session were computed (*prctile* function in Matlab).
5. Pairwise activity correlations between cells (each session). Activity trajectories were compared to each other using pairwise Spearman correlations. These correlations were performed between bins of cells as well as between individual cells within defined cell groups (MALE and OTHER cells), as described below.

##### Bins of cells (Figure 3C)

In each session, cells were sorted by their separability values and binned in 2.5 percentile-width bins. For example, the first bin consisted of cells with separability values between the 0^th^ (minimum) and 2.5^th^ percentile of the distribution for that session. Then, to obtain a representative activity trajectory for each bin on a given trial, the activity of each cell in the bin was taken over timepoints and summarized over cells (median). This gave one activity vector per bin on a given trial.

Activity vectors were then pairwise correlated between bins, giving a bins x bins matrix of correlation coefficients (Spearman *ρ*) on a given trial. This matrix was summarized (median) over trials of a given type in the session (e.g., male trials), giving a pairwise correlation matrix for that trial type. If the session had no cells in one or more bins, the matrix values associated with those bins were set to NaN and ignored in the subsequent all-session summary (see below).

The correlation matrix for a given trial type (e.g., male trials) was summarized over sessions (median) in each experimental group to obtain a representative correlation matrix for that group (as shown in Figures 3D, F; S7C, E).

##### Individual cells within cell groups

Within a given cell group (MALE or OTHER cells), the activity of each cell was taken over timepoints on a given trial. This activity vector was then correlated to the vectors of all other cells in the group to obtain a set of correlation coefficients (Spearman *ρ* values). These coefficients were summarized over all unique cell pairs (median) to obtain one value per trial. Values were then summarized over all trials of a given type in the session (e.g., median of male trials). This gave 4 summary *ρ* values for each session, corresponding to the 2 cell groups and 2 trial types: MALE cells on male trials, OTHER cells on male trials, MALE cells on milk trials, and OTHER cells on milk trials (as shown in Figures 3E, G; S7D, F).

6) Definition of MALE and OTHER cell groups (each session). MALE cells were detected from the pairwise correlation matrix in Figure 3D (on male trials in control group) using change point analysis (Figure S7A). The upper-right triangle of the correlation matrix was considered. The coefficients in each row were summarized (median), giving a vector of values over bins. The vector was then checked for outliers (Grubbs’ test), which could obscure the detection of a stable change point. If there were no outliers, the residual sum of squares (RSS) between the data values and the overall mean was calculated (RSS0). Then, a candidate change point was identified as a break point that maximally reduced the RSS compared to RSS0 (best improvement). The RSS of the candidate change point (RSS1) was calculated using the data values and two segment means defined by the break (see horizontal lines). To test for significance, the observed best-improvement value (RSS1-RSS0) was compared to a permutation distribution of best-improvement values generated by randomizing cell order and recomputing the correlation matrix in Figure 3D (*n* = 1000 iterations, one-sided test with α = 0.05). If the observed best-improvement value was significant, MALE cells were defined as the group of cells with separability values at or above the change point. OTHER cells were defined as the group of all remaining cells (separability below change point).

When testing for milk-representing cells (Figure S7G), the lower-left triangle of the correlation matrix in Figure S7C was used. The same approach as described above was applied.

### Computational modeling

Model simulations were implemented using custom code in Python (version 3.9.13).

#### Model description

The model consisted of a population of pyramidal neurons and a population of interneurons (OXTR neurons). A given pyramidal neuron fired at time *t* with a probability set by a Bernoulli distribution (*B*). The input to *B* depended on the activity of other pyramidal neurons and the interneuron population, as follows:

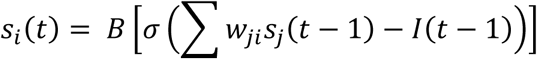

*s_i_(t)* represented the firing state of neuron *i* (0 for non-firing, 1 for firing), *σ* a rescaled sigmoid bounded between 0.1 and 0.4, and *w_ji_* the weight of the connection from pyramidal neuron *j* to neuron *i*. *I(t)* was a global inhibitory input from OXTR neurons, whose strength at time *t* was proportional to the sum of all pyramidal firing events, as follows:

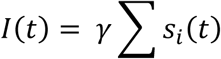

The pyramidal population was split into two groups (MALE and OTHER cells), which had distinct connectivity. All MALE cells were connected to each another with the same weight *w_M_*, and all the OTHER cells were connected to each other with the same weight *w_O_*, resembling a mean field model. There were no connections between the two groups.

This architecture gave the parameters *w_M_ = s/N_M_, w_O_ = s/N_O_, γ = q/N*, where *N_M_* and *N_O_* were the numbers of MALE and OTHER cells. *N* was the total number of pyramidal neurons (MALE + OTHER). *s* and *q* were scaling parameters, where *q* set the strength of inhibition.

#### Parameter selection

Considering the expected firing rate of a neuron in either the MALE and OTHER cell groups at time *t* (*a_M_(t)* and *a_O_(t)*), the system dynamics could be written for large *N* as follows:

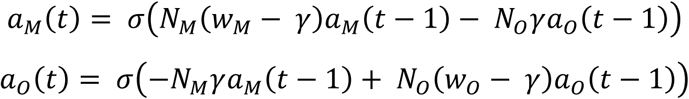

At equilibrium, the expected firing rates would not change over time (i.e., *a_M_(t) = a_M_(t-1)* and *a_O_(t) = a_O_(t-1)*). Setting both groups’ firing rates to be equivalent at equilibrium (*a_O_(t) = a_M_(t)*) led to *N_M_w_M_ = N_O_w_O_*. This equality specified the weights *w_M_* and *w_O_*, which grew inversely to the number of neurons in the corresponding group. The numbers of neurons were taken to be *N_M_* = 20 and *N_O_ =* 80 for the main model, and *N_M_* = 50 and *N_O_ =* 50 for the symmetric model.

In the reference (OXTR ON) condition, the scaling parameters *s* and *q* were manually chosen to (1) set the expected firing rate at equilibrium to be low (*q* > *s*) and (2) ensure that fluctuations in the expected firing rate around the equilibrium point would be sustained for many time steps but still remain close to the equilibrium point. *s* and *q* were thus selected to be 10 and 15, which produced an equilibrium point of 0.16. This value was used as the baseline firing rate in simulations.

In the OXTR OFF condition, *q* was reduced from 15 to 1.5 such that pyramidal activity was not controlled by inhibition. Reducing *q* allowed the expected pyramidal firing rate to strongly deviate from the equilibrium point. Because of the absence of inhibition and the strong connectivity within groups, the expected firing rate of MALE and OTHER cells would be self-reinforcing until both groups were close to the maximum value allowed by *σ* (0.4).

#### Simulations of neural activity and behavioral choice

Neural activity was simulated on 500 OXTR ON trials and 500 OXTR OFF trials. On each trial, firing was simulated over 120 time steps, starting at a baseline firing rate. Each neuron’s firing events were summed over the last 20 steps and the rate computed (number of firing events / 20 time steps). This rate was summarized (median) over all neurons in a group (e.g. MALE cells) to obtain the group’s observed firing rate (Figures 4B, F).

To make a behavioral choice (male or milk) on each trial based on the relative firing rates of MALE versus OTHER cells, a voting process was used. This process assumed that a given neuron from each group had the same voting power, such that one MALE cell firing at a specific rate would compensate an OTHER cell firing at the same rate. Thus, to take a decision, the model compared the activity of both groups, first adding the firing rates over cells within each group individually and then taking the difference between groups. The difference was then multiplied by 3 to increase the sharpness of the decision boundary. This value was fed into a sigmoidal function biased to be 0.5 at the equilibrium point and rescaled to be bounded between 0.1 and 0.9. The output was used as the probability of a Bernoulli trial determining the behavioral outcome as male or milk.

To obtain representative summary measures (see next section), multiple simulation runs (*n* = 12) were performed. This gave 12 sets of 500 OXTR ON trials and 500 OXTR OFF trials.

#### Comparison of simulated OXTR ON and OFF trials

For a given set of 500 trials (OXTR ON or OFF), male preference was calculated as the proportion of male trials (see “Male and female preference” section above). The ability of MALE cells to distinguish between male and milk trials was quantified as the AUC (taking x-axis values of Figure 4B for the main model and Figure 4F for the symmetric model). Male preference and MALE AUC values were then summarized (median) over simulation runs.

#### Predicted consistency with experimental data

Given the equality of voting power across neurons, the choices of the main (asymmetric) model would be expected to be milk-biased when both populations were firing close to their maximum value (i.e., in the OXTR OFF condition).

Furthermore, the two cell groups differed in the number of neurons, and thus also in the strength of their self-reinforcing weights (according to *N_O_w_O_ = N_M_w_M_*). MALE cells had stronger self-reinforcing weights. Deviations from equilibrium would thus be expected to more strongly propagate within the MALE cell group over time steps on a given trial, leading to a greater activity range over trials.

### Statistics

Unless otherwise indicated, data are shown as median and interquartile range (IQR). Boxplot whiskers extend to the most extreme data points within the range of *Q1* - 1.5 x (*Q3* - *Q1*) and *Q3* + 1.5 x (*Q3* - *Q1*), where *Q1* and *Q3* represent the 25^th^ and 75^th^ percentiles, respectively.

Statistical testing was performed in RStudio (version 2025.09.2; R version: 4.5.2) and Matlab. Linear mixed-effects models (LMEs) and binomial generalized linear mixed models (GLMMs) were applied to data where measurements were repeated within subjects (e.g., multiple sessions per subject)^60^. These models explicitly accounted for within-subject dependencies by including subject id as a random effect. Binomial GLMMs (with logit link function) were used when the outcome variable was a proportion based on counts (e.g., male or female preference, which were based on social and milk trial counts). The *lme4* and *lmerTest* packages and *anova* function were used to fit and evaluate the models. The *emmeans* package was used for post hoc tests. When applying model-based t-tests or a Type III ANOVA on a model, the degrees of freedom were estimated using the Kenward-Roger method. In model summaries, *t* refers to the t-test, *Z* refers to the Wald Z-test, and *F* refers to the ANOVA F-test.

In all applicable cases, two-sided tests were used. The only exceptions were the Two One-Sided Tests procedure (see below), permutation testing to define MALE cells (see “Definition of MALE and OTHER cell groups” above), and the analysis of whether estrous state modulated the difference between actual and predicted male preference (Figure 1H). For this analysis, previous work^22^ suggested a directionality of the effect (SR lower than NR), motivating a one-sided test.

Multiple comparisons were corrected using the Benjamini-Hochberg (false discovery rate: 0.05) or Holm procedures, as specified. In labeling statistical results, * indicates *P* < 0.05, ** *P* < 0.01, and *** *P* < 0.001.

#### Statistical methods specific to Figures 1E-H, S5B

In each experimental group (control, Halo), a binomial GLMM was fit to male preference values (underlying trial counts) in pre-opto. sessions, using session-matched female preference (mean-centered) as a fixed effect and subject ID as a random intercept. However, in both groups’ models, the contribution of the random intercept was negligible (very small estimated variance of subject ID), producing a singular fit warning. Therefore, both control and Halo models were simplified to standard binomial logistic regression models (outcome: male preference (underlying trial counts); fixed effect: mean-centered female preference; no random effect).

Each group’s logistic regression model (slope and intercept) was then used to predict male preference in opto. sessions (Figures 1E-H) or pre-opto. sessions (Figure S5B) based on the observed female preference. This gave a set of predicted and actual (observed) male preference values. To test for a difference between actual and predicted values, a binomial GLMM was fit to this data. This model estimated a coefficient for the difference between actual and predicted values (logit scale) in each experimental group. A significant group coefficient indicated a meaningful difference.

In addition, the Two One-Sided Tests (TOST) procedure was used to test whether actual and predicted male preference were equivalent to each other within a specified tolerance range (i.e., model-estimated deviation lies within range).

To specify the tolerance, it was assumed that the number of male trials in a session could vary by +/- 1 by chance. Using the median predicted preference as a reference level, +/- 1 male trials corresponded to a preference boost or loss of 1/16 relative to this reference, where 16 was the typical number of available trials in a session. The maximum magnitude of this boost/loss on the logit scale was used as the tolerance. Two one-sided Wald Z-tests were used to test whether the model-estimated deviation between actual and predicted male preference fell within both ends of the tolerance range (+/- tolerance value). The maximum *P* value of the two tests was reported.

## Supporting information

Legends for Videos S1 & S2

## Data availability

The data collected and analyzed in this study are available from the corresponding authors upon reasonable request.

## Code availability

Custom code written for data analysis in this study are available from the corresponding authors upon reasonable request.

## Acknowledgements

We thank the UZH LASC team for excellent animal husbandry, Simone Holler for histology assistance, Giorgio Rizzi and Harald Dermutz for technical assistance, and Valerio Mante for valuable feedback and discussions. This work was supported by the Swiss National Science Foundation (CRSII5-173721 and 315230 189251 to B.F.G.), ETH Zurich project funding (ETH-032 24-2 to B.F.G.), and Human Frontier Science Program (LT000539/2018 to E.A.A.).

## Author contributions

E.A.A. and B.F.G. conceived the project. E.A.A., R.B., and R.L. designed and performed mouse experiments, including population calcium imaging (E.A.A., R.B.) and patch clamp electrophysiology (R.L.). R.B. managed the breeding and histology of experimental animals. R.B. and E.A.A. performed stereotactic surgeries and baseplate mounting, respectively. E.A.A. and E.S.M. analyzed population calcium imaging data with conceptual and technical support from B.E. R.L. analyzed patch clamp electrophysiology data. P.V.A. conceived and implemented the computational model with input from E.A.A. E.A.A., P.V.A., R.L., and R.B. wrote the paper with input from all of the authors.

## Declaration of interests

The authors declare no competing interests.

## Materials & Correspondence

Correspondence and material requests should be addressed to Elizabeth A. Amadei and Benjamin F. Grewe.

---

**Figure S1.**
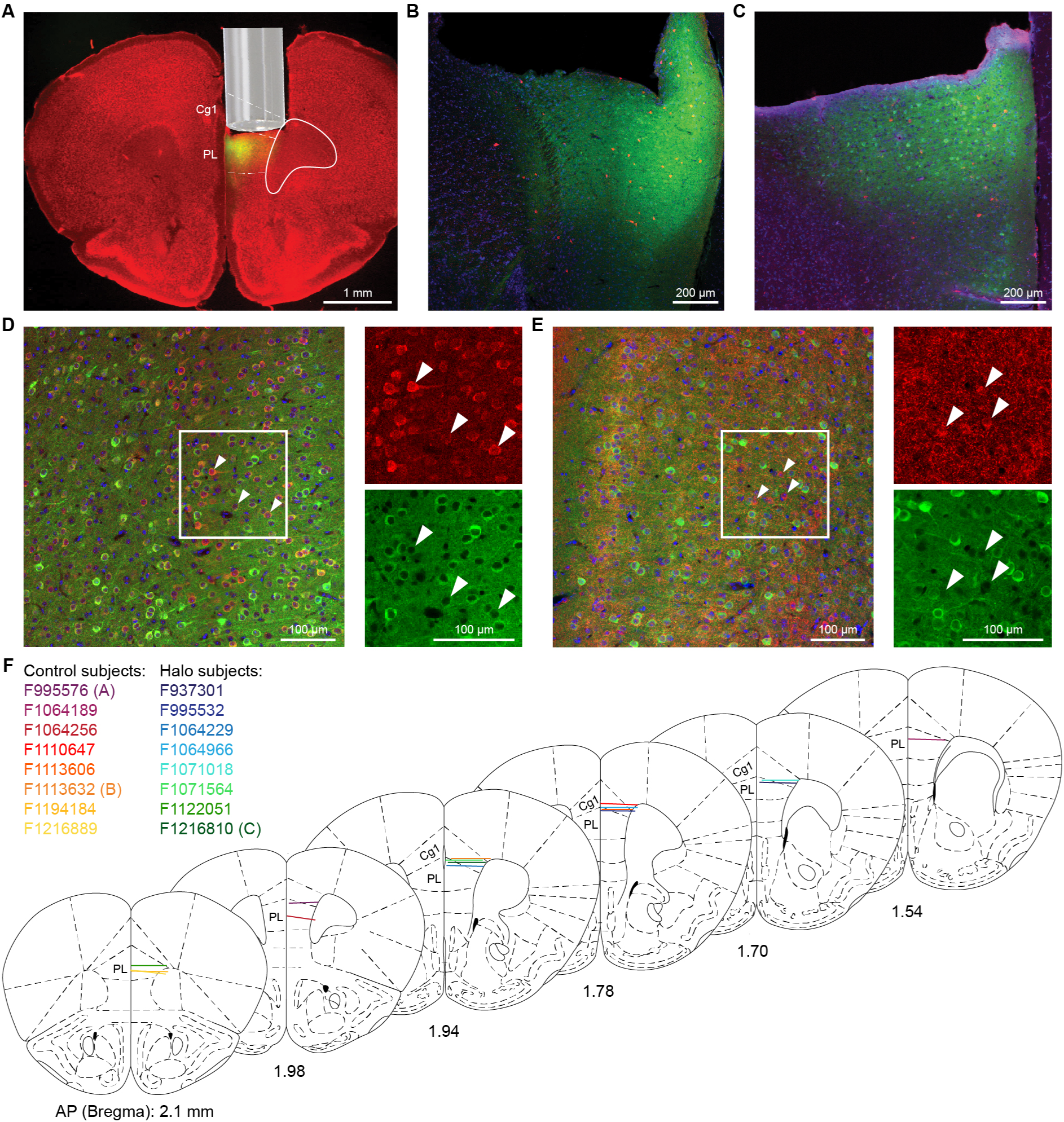
Validation of viral expression and implant placements in mPFC. (**A**) Example viral expression and implant location in the right mPFC of subject F995576 (implant locations of all subjects shown in (**F**)). GCaMP6m expression (green) and Nissl stain (red). PL: prelimbic cortex. Cg1: cingulate cortex, area 1. (**B**) Example GCaMP6m (green) and mCherry (red) expression in control subject F1113632. DAPI stain (blue). (**C**) Example GCaMP6m (green) and Halorhodopsin-mCherry (red) expression in Halo subject F1216810. DAPI stain (blue). (**D-E**) GCaMP6m expression occurs in Neurogranin-positive cells (neural marker) but not glutamate decarboxylase (GAD)-positive cells (GABAergic marker), consistent with pyramidal expression. (**D**) GCaMP6m (green), Neurogranin (red) and DAPI (blue). Right top: Zoom-in on Neurogranin stain; Right bottom: Zoom-in on GCaMP6m expression. Arrows point to three example Neurogranin-positive neurons expressing GCaMP6m. (**E**) GCaMP6m (green), GAD (red) and DAPI (blue). Right top: Zoom-in on GAD stain; Right bottom: Zoom-in on GCaMP6m expression. Arrows point to three example GAD-positive neurons that do not express GCaMP6m. (**F**) Locations of the implant base for all subjects (*n* = 8 control, 8 Halo). Letters in parentheses (e.g., “(A)”) indicate figure panels showing representative images from these subjects.

**Figure S2.**
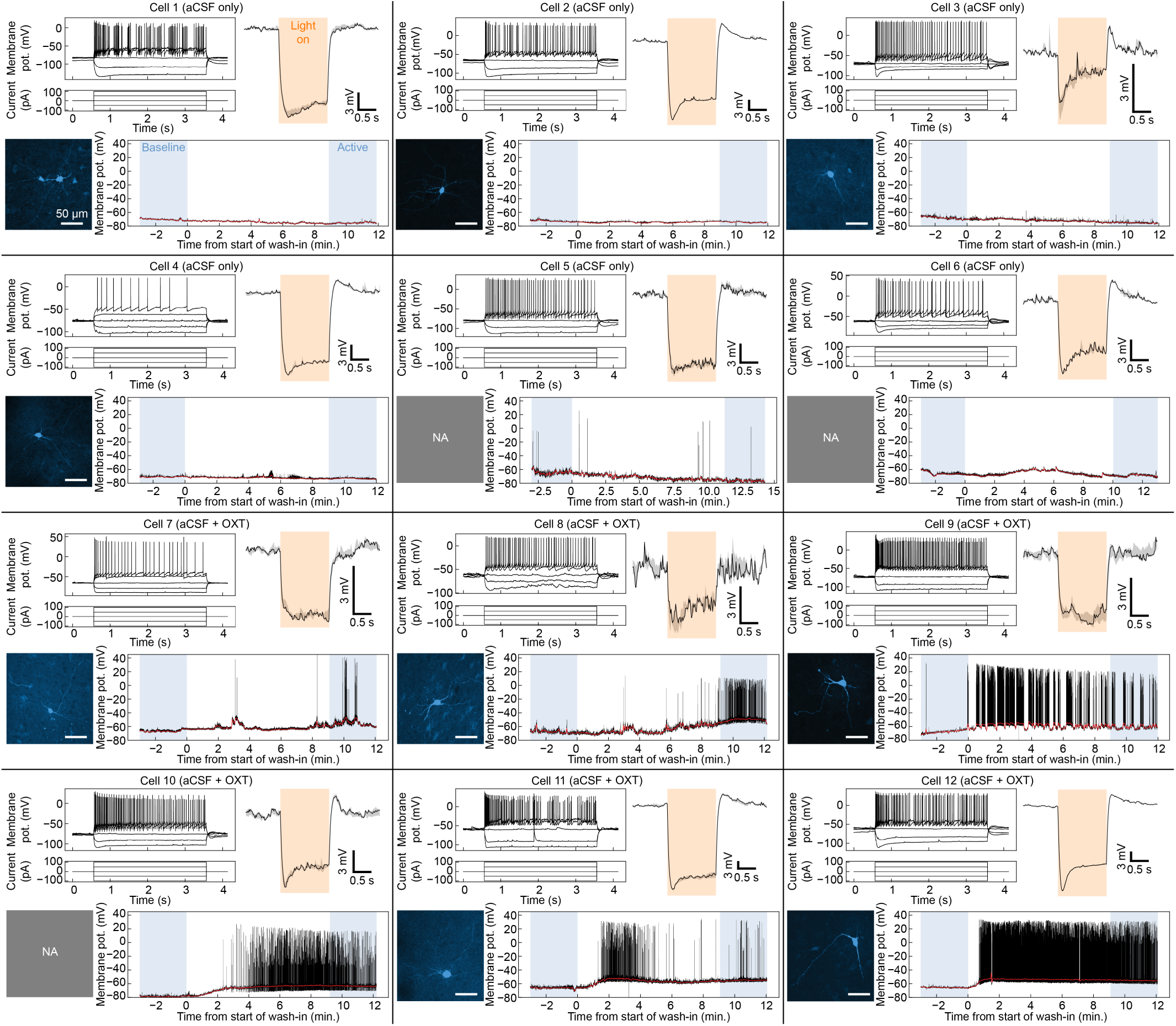
Validation of Halorhodopsin-expressing mPFC OXTR neurons. Current clamp recordings of Halorhodopsin-expressing mPFC OXTR neurons during wash-in experiments with aCSF only (*n* = 6 cells) or aCSF + oxytocin (OXT; *n* = 12 cells; 6 shown here and remaining 6 shown in Figure S3) for at least 12 minutes. Each panel shows one neuron and includes: Top left: Membrane potential response to negative and positive current injections (50 pA steps, 3 s). Top right: Validation of inhibitory response to 1.5 s optogenetic light exposure. Thick line and shaded area show mean +/- standard error of the mean (SEM) over 5 repetitions. Bottom left: Histological reconstruction of recorded neuron using biocytin-streptavidin staining. All reconstructed neurons display a stellate-like morphology, consistent with previous work identifying OXTR neurons in mPFC^22^. No reconstruction available for cells 5, 6, and 10. Bottom right: Raw (black) and low-pass filtered (red) recordings during wash-in experiments. Wash-in began at time 0 and lasted until the last timepoint. Blue shaded areas indicate a “baseline window” (180 s before wash-in starts) and an “active window” (last 180 s of wash-in), used for quantifications.

**Figure S3.**
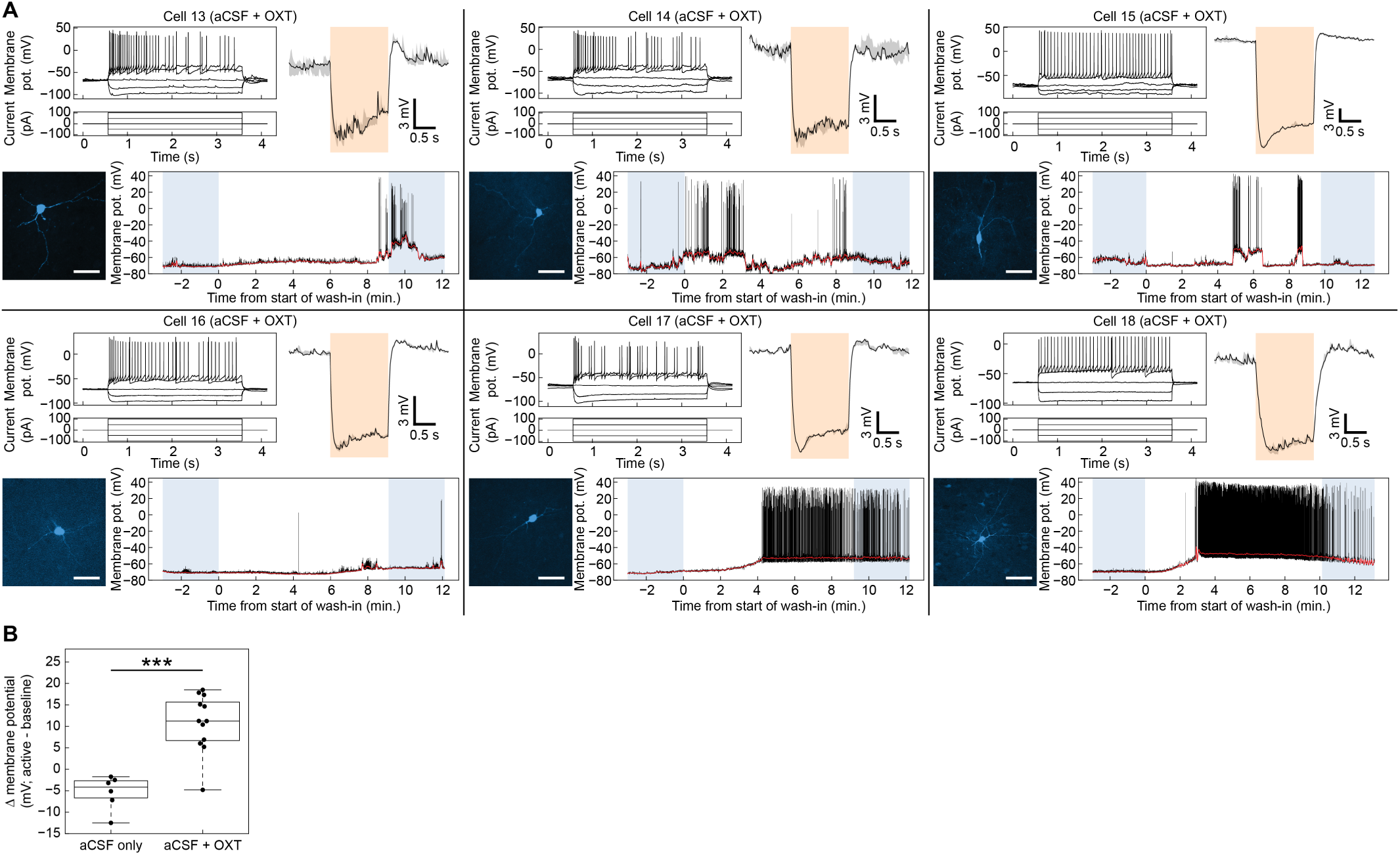
Validation of Halorhodopsin-expressing mPFC OXTR neurons (Continued). (**A**) Continued from Figure S2: remaining 6 Halorhodopsin-expressing mPFC OXTR neurons receiving aCSF + OXT wash-in. (**B**) Change in membrane potential from baseline to active windows for all neurons exposed to aCSF only (*n* = 6) or aCSF + OXT (*n* = 12). For each neuron, the membrane potential was low-pass filtered and averaged over all timepoints within each window to produce one baseline value and one active value. Then the change from baseline to active was calculated. OXT exposure significantly increased the membrane potential compared to aCSF only (Wilcoxon rank-sum test, *P* < 0.001***). Boxplots show median and IQR over neurons.

**Figure S4.**
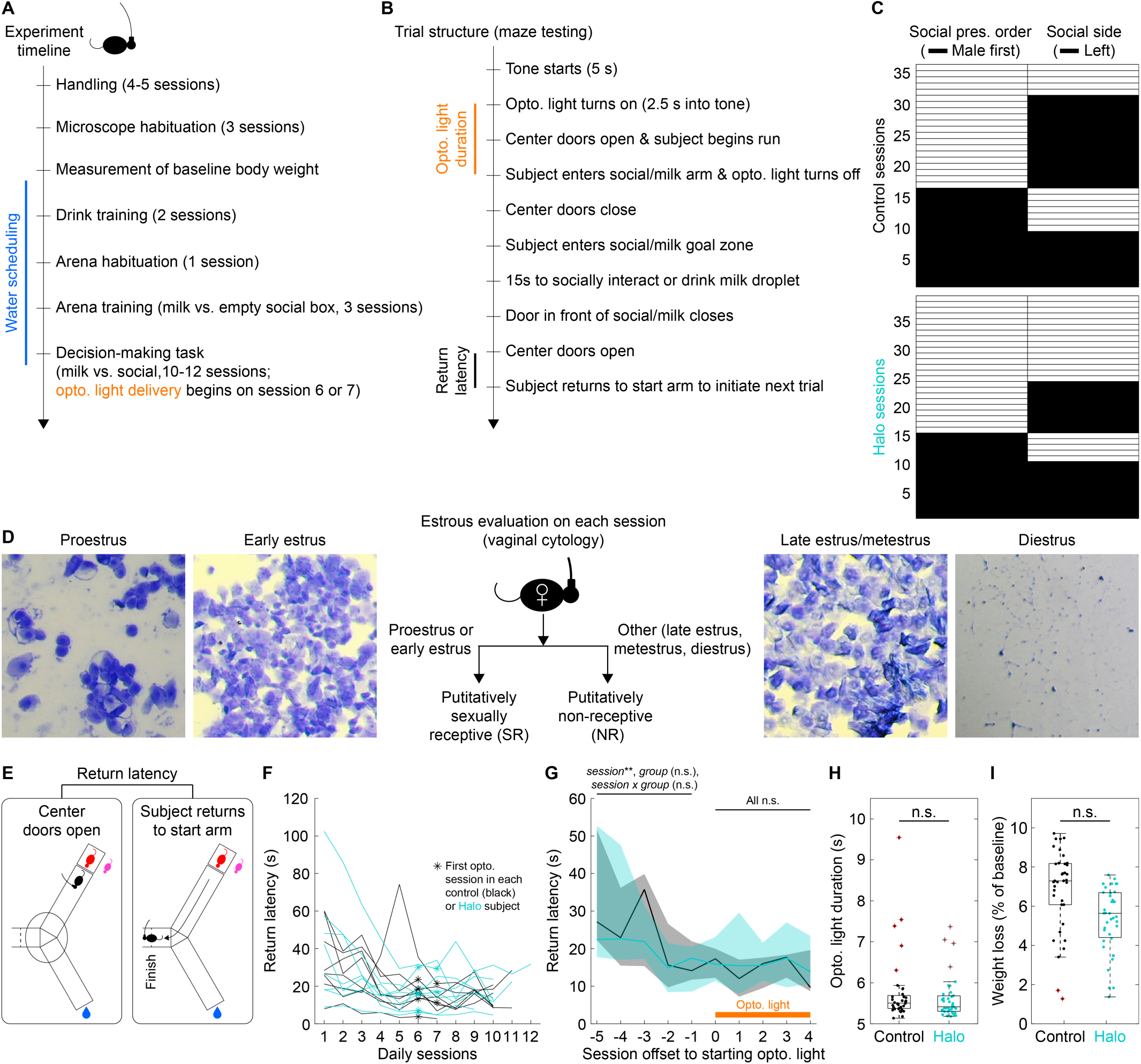
Experimental design. (**A**) Experiment timeline for each subject. All sessions were performed daily, with the exception of arena habituation, which occurred at the end of the second day of drink training. (**B**) Trial structure, including definitions of optogenetic light duration and return latency. (**C**) Distribution of social stimulus presentation order (male or female first) and location (left- or right-hand side of maze relative to start arm) over optogenetics sessions used in behavioral and neural analyses (*n* = 36 control, 39 Halo sessions). Sessions sorted by presentation order and social side. (**D**) Representative crystal violet-stained vaginal samples taken at the end of each session. Based on estrous phase, sessions were classified as putatively sexually receptive or non-receptive. (**E**) Definition of latency to return to start arm at the end of each trial, used as a measure of task performance (lower values indicate better task performance). (**F**) Summarized return latencies in individual control (*n* = 8) and Halo (*n* = 8) subjects over maze testing sessions (median over all trials in each session). Optogenetic light was introduced on session 6 or 7, once a subject’s return latency had been assessed to be stable (see verification in (**G**)). (**G**) Same data as (**F**) but now aligning sessions to the start of optogenetic light (5 sessions before and 5 sessions with optogenetic light; *n* = 7-8 control and 7-8 Halo subjects at each session index). Thick lines and shaded areas show median and IQR over subjects in each group. During pre-opto. sessions, there was a significant effect of session on return latency, but no significant effect of group (control, Halo) or session x group interaction (LME with session (mean-centered) and group as fixed effects: *F_session*(1, 14) = 10.02, *P_session* = 0.007**; *F_group*(1, 14) = 0.03, *P_group* = 0.862; *F_interaction*(1, 14) = 0.39, *P_interaction* = 0.544). This indicated that return latency improved over pre-opto. sessions, with a similar trajectory in control and Halo groups. During opto. sessions, there were no significant effects (*F_session*(1, 58.74) = 0.66, *P_session* = 0.419; *F_group*(1, 13.97) = 0.80, *P_group* = 0.386; *F_interaction*(1, 58.74) = 0.14, *P_interaction* = 0.707), indicating that the return latency was stable and similar between groups. (**H**) Duration of optogenetic light in control versus Halo sessions (same *n* as (**C**); light duration summarized over all trials within each session to obtain one value per session). Halo group did not differ from control (LME with group as fixed effect: *Estimate* = -0.127, *SE* = 0.309, *t*(13.99) = -0.41, *P* = 0.686). (**I**) Weight loss in control versus Halo opto. sessions (same *n* as (**C**)). Weight loss computed relative to a baseline measurement before water scheduling (see (**A**)). Halo group did not differ from control (LME with group as fixed effect: *Estimate* = -1.413, *SE* = 0.871, *t*(13.99) = -1.62, *P* = 0.127). Boxplots in (**H-I**) show median and IQR over sessions.

**Figure S5.**
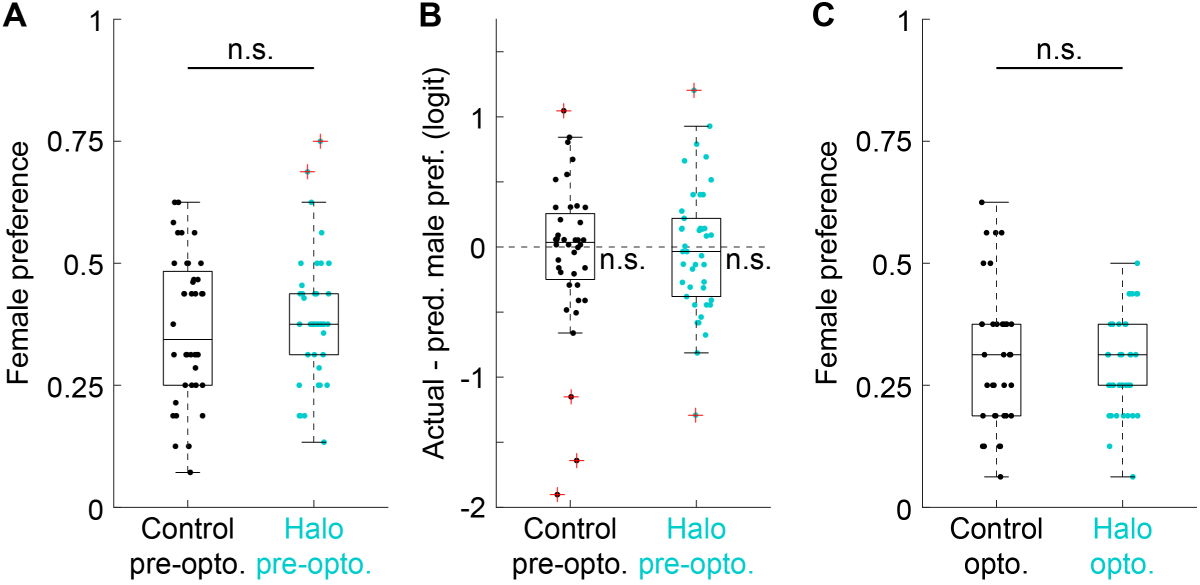
Binomial logistic regression adequately fits pre-opto. sessions, and female preference does not differ between control and Halo groups. (**A**) Comparison of female preference between control (*n* = 40) and Halo (*n* = 42) sessions before optogenetic light stimulation (pre-opto. sessions; binomial GLMM with group as fixed effect: *Estimate* = 0.119, *SE* = 0.246, *Z* = 0.49, *P* = 0.628). (**B**) Actual - predicted male preference in control and Halo pre-opto. sessions (same *n* as (**A**); predictions for both groups from pre-opto. regression model fits in Figure 1E). Session values transformed to logit scale to match binomial GLMM used for statistical testing. No significant difference between actual and predicted male preference in either group (control: *Estimate* = 0.000, *SE* = 0.086, *Z* = 0, *P* = 1; Halo: *Estimate* = 0.000, *SE* = 0.083, *Z* = 0, *P* = 1). In both groups, actual and predicted male preference were significantly equivalent within a tolerance range of +/- 1 male trials (TOST: *P control* and *P Halo* both < 0.001***). (**C**) Comparison of female preference between control (*n* = 36) and Halo (*n* = 39) opto. sessions (binomial GLMM with group as fixed effect: *Estimate* = -0.027, *SE* = 0.246, *Z* = -0.11, *P* = 0.914). Throughout the figure, boxplots show median and IQR over sessions.

**Figure S6.**
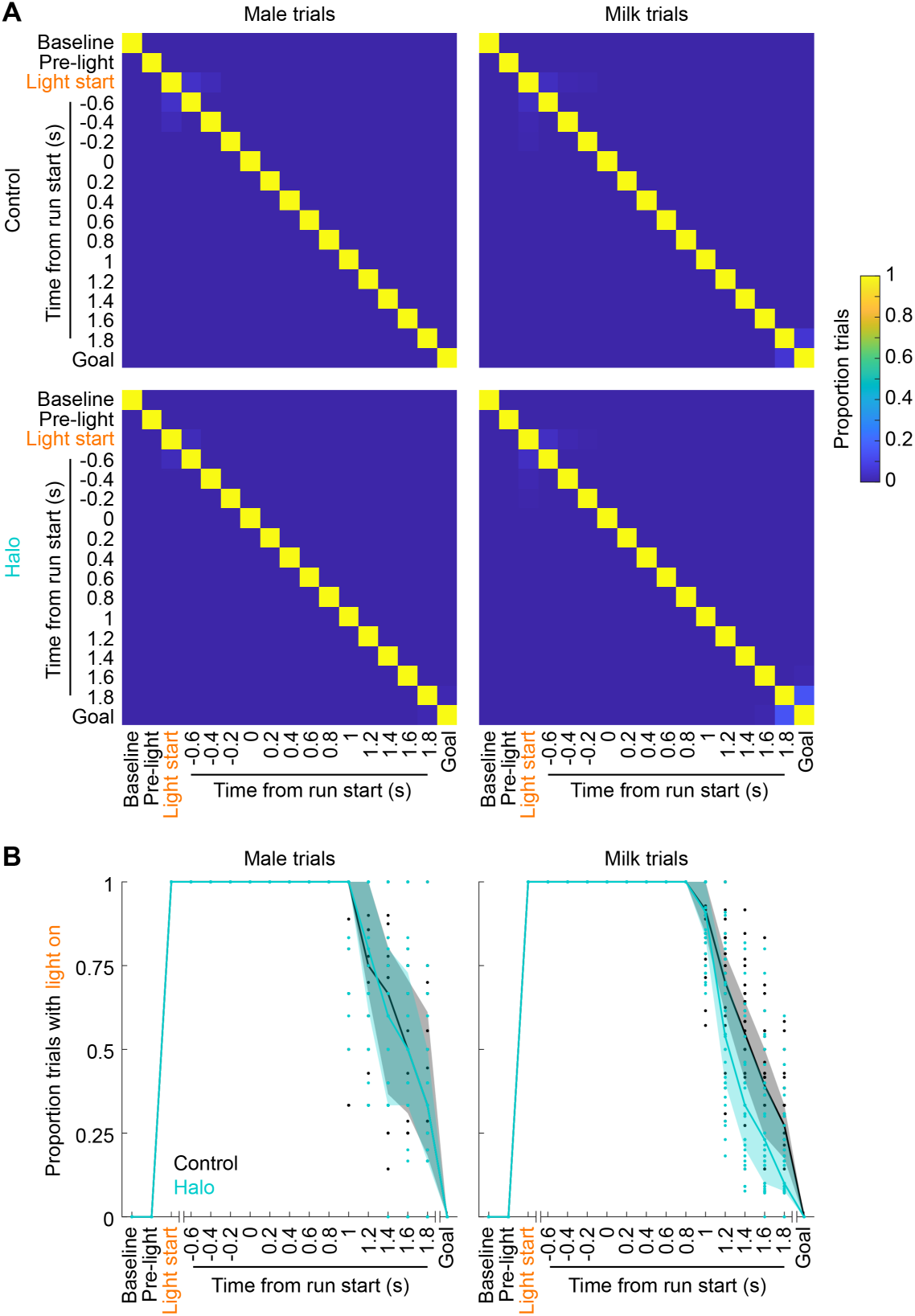
Characterization of trial timepoints in relation to each other and optogenetic light. (**A**) Overlap between timepoints on male and milk trials (columns) in control and Halo groups (rows). Within a given plot, each row and column indicate one timepoint. For each row, the proportion of trials where a given timepoint overlaps with itself and each of the other timepoints is indicated by a color from dark blue (0) to yellow (1). In control group, *n* = 173 male and 402 milk trials (pooled over opto. sessions). In Halo group, *n* = 162 male and 461 milk trials. Deviation from dark blue outside of the diagonal shows the extent of overlap between timepoints. This deviation was low across trial types and groups, indicating that timepoints were largely and consistently distinct. (**B**) At each timepoint, proportion of trials of a given type (male or milk) containing optogenetic light. Proportions were computed on trials from each session (*n* = 36 control sessions and 39 Halo sessions). Dots show individual sessions. Thick lines and shaded areas show median and IQR over sessions in each group.

**Figure S7.**
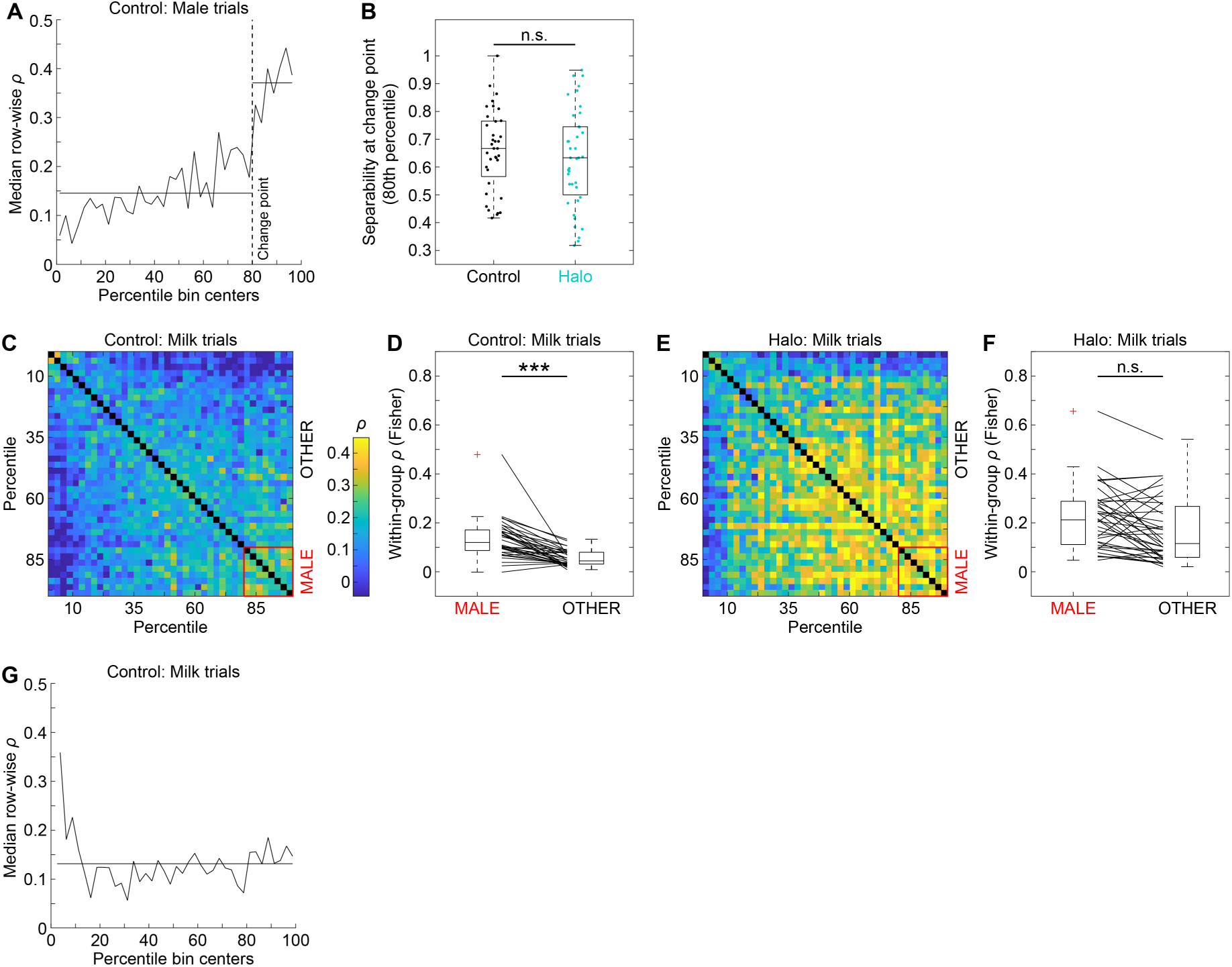
Detection and characterization of MALE cells. (**A**) To detect MALE cells, the upper-right triangle of the correlation matrix in Figure 3D was taken. Each row was summarized (median, excluding diagonal), giving a vector of correlation coefficients plotted here. The vector was then checked for outliers (Grubbs’ test), which could obscure the detection of a stable change point. If there were no outliers, the residual sum of squares (RSS) between the data values and the overall mean was calculated (RSS0). Then, a candidate change point was identified as a break point that maximally reduced the RSS compared to RSS0 (best improvement). The RSS of the candidate change point (RSS1) was calculated using the data values and two segment means defined by the break (see horizontal lines). To test for significance, the observed best-improvement value (RSS1-RSS0) was compared to a permutation distribution of best-improvement values generated by randomizing cell order and recomputing the correlation matrix (*n* = 1000 iterations; see Methods). This approach gave a significant (*P* < 0.001***) change point at the 80^th^ percentile of the separability distribution (vertical dotted line). (**B**) Separability at the 80^th^ percentile of each session’s separability distribution (*n* = 36 control, 39 Halo sessions). The Halo group showed comparable separability values to control (LME with group as fixed effect: *Estimate* = -0.036, *SE* = 0.068, *t*(13.97) = -0.53, *P* = 0.605). (**C**) Same summary correlation matrix as Figure 3D but now computed from cells’ activity on milk trials (as opposed to male trials) in control sessions. (**D**) In control sessions (*n* = 36, individual lines), MALE cells showed more strongly correlated activity compared to OTHER cells (LME with MALE - OTHER coefficients as outcome variable: *Estimate* = 0.079, *SE* = 0.014, *t*(6.66) = 5.59, *P* < 0.001***). Correlation coefficients were Fisher-transformed for statistical testing. (**E-F**) Same as (**C-D**) but now in Halo sessions (*n* = 39). MALE and OTHER cells did not differ in their strength of correlated activity in Halo sessions (LME with MALE - OTHER coefficients as outcome variable: *Estimate* = 0.049, *SE* = 0.025, *t*(6.99) = 1.95, *P* = 0.093). (**G**) To detect any milk-representing cells, the lower-left triangle of the correlation matrix in (**C**) was taken. As in (**A**), each row was summarized (median, excluding diagonal), giving a vector of correlation coefficients plotted here. A break between the first and remaining values maximally reduced the RSS. However, the first value was an outlier (Grubbs’ test), which prevented considering this break as a change point. Horizontal line shows mean over all values. In (**B, D, F**), boxplots show median and IQR over sessions.

## Notes

### Competing Interest Statement

The authors have declared no competing interest.

### Summary of Updates

Updated legends for Videos S1 & S2

