## Supplementary material for "Inhibition tunes prefrontal circuit dynamics to promote sociosexual behavior in female mice": Legends for Videos S1 & S2

**Video S1 | Example social trial.** Side-by-side video of behavioral recording (left) and mPFC population calcium imaging (right; ∆F/F) in an example Halo subject (F1064229) during an example male trial. Left: White box indicates tone initiating trial start. Orange box indicates optogenetic light. Right: scale bar is 100 µm.

**Video S2 | Example milk trial.** Side-by-side video of behavioral recording (left) and mPFC population calcium imaging (right; ∆F/F) in an example Halo subject (F1064229) during an example milk trial. Left: White box indicates tone initiating trial start. Orange box indicates optogenetic light. Right: scale bar is 100 µm.
